# Factorial study of the RNA-seq computational workflow identifies biases as technical gene signatures

**DOI:** 10.1101/2020.01.30.924092

**Authors:** Joël Simoneau, Ryan Gosselin, Michelle S. Scott

## Abstract

RNA-seq is a modular experimental and computational approach that aims in identifying and quantifying RNA molecules. The modularity of the RNA-seq technology enables adaptation of the protocol to develop new ways to explore RNA biology, but this modularity also brings forth the importance of methodological thoroughness. Liberty of approach comes with the responsibility of choices, and such choices must be informed. Here, we present an approach that identifies gene group specific quantification biases in currently used RNA-seq software and references by processing sequenced datasets using a wide variety of RNA-seq computational pipelined, and by decomposing these expression datasets using an independent component analysis matrix factorisation method. By exploring the RNA-seq pipeline using a systemic approach, we highlight the yet inadequately characterized central importance of genome annotations in quantification results. We also show that the different choices in RNA-seq methodology are not independent, through interactions between genome annotations and quantification software. Genes were mainly found to be affected by differences in their sequence, by overlapping genes and genes with similar sequence. Our approach offers an explanation for the observed biases by identifying the common features used differently by the software and references, therefore providing leads for the betterment of RNA-seq methodology.

## INTRODUCTION

Modularity is both a boon and a burden for RNA-sequencing (RNA-seq) analysis. At its core, RNA-seq leads to the identification and quantification of RNA molecules from a biological extract^1^. RNA-seq is in fact an umbrella term, encompassing a broad diversity of laboratory and computational design choices, where each choice defines the scope of the study, the questions it might answer^2^. The modularity of RNA-seq has been a steppingstone in the development of many other techniques, mainly differing by the way in which the RNA is extracted, and with consequent modifications of the *in silico* pipeline. For example, ribosome profiling (Ribo-seq) can be summarized as the RNA-seq of the RNA fragments protected by ribosome footprints^3^. The modularity of the RNA-seq *in silico* pipeline, stemming from the usage of well-defined data processing steps, each supported by specific file formats, has led to the creation of specialized software for each of the different steps, compartmentalizing the data processing. Having defined data processing steps helps to isolate each technical problem, creating an ecosystem where research groups specialize in answering individual steps, and proceed to benchmark accordingly.

Due to the RNA-seq modular nature, one has to constrain many degrees of freedom linked to the experimental design to be able to generate and process the data. These degrees of freedom represent protocols, reagents and kits on the experimental side, and software, references and parameters on the computational side. With a benchmarking point of view, experimental and computational design choices mainly differ by their permanence. While a specific RNA extract can only be processed once in the laboratory, sequencing being a destructive method, sequencing data can theoretically be reanalyzed in a wide variety of ways. This facilitates the creation of large-scale benchmarking of the RNA-seq *in silico* pipeline, because they can be built on a very specific and unique group of datasets, while benchmarking studies of the RNA-seq laboratory pipeline have to deal with extra noise and reproducibility issues coming from biological variability. RNA-seq in silico pipeline benchmarking studies are also cheaper to produce since they only require computational resources.

Benchmarking the RNA-seq pipeline may take two different approaches, either analytical or systemic. In an analytical approach, the pipeline would be studied in its irreducible form, meaning that each step would be benchmarked independently. The analytical approach is usually used when publishing a new tool. To ensure the relevance of the newly proposed method, authors will compare its performance to current methods^4^. Depending on the position of the studied step in the overall pipeline, it might be difficult to meaningfully assess its quality. For example, alignment software has often been characterized using the percentage of alignment as a metric to be optimized, even though such a metric does not hold any biological meaning^5^. To bypass the need for other metrics, it is also possible to study the effects of a given step on the rest of the pipeline, using a fixed downstream processing. Conversely, in a systemic approach, the pipeline is studied as a whole. While the analytical approach is based on the hypothesis that each step is fully independent, the systemic approach can be used to study interactions between steps.

In Table 1, we compiled a list of articles benchmarking the RNA-seq in silico pipeline by considering more than one step in their analysis^6–15^. The main point of interest is the imbalance between the pipeline steps, both in the number of elements studied per article as well as globally. These articles often use the term “RNA-seq workflow” to describe the object of their analysis, while also mainly limiting themselves to the alignment and quantification software. As we described earlier, the RNA-seq *in silico* workflow should consider all steps from the raw FASTQ files to the count matrices^16^. Insofar as we do not have any study highlighting the importance, or lack of, of every workflow step, overlooking some steps might hide important biases. In a previous study, we highlighted that the trimming step and the choice of genomic annotations were often not reported in methodologies of articles performing RNA-seq^17^. Furthermore, we can observe that these steps are also overlooked in the articles studying the RNA-seq workflow (Table 1). Only one article included in our analysis reported using more than one annotation reference, and only used it to evaluate transcript assembly. Many of those benchmarks also did not provide any information about trimming. We now find ourselves before a circular causality problem in which we are not benchmarking certain steps because they are not being reported and we are not reporting them because they are not being benchmarked. In any case, there is insufficient data for a meaningful answer regarding whether the overlooked information holds any importance.

**Table 1.**
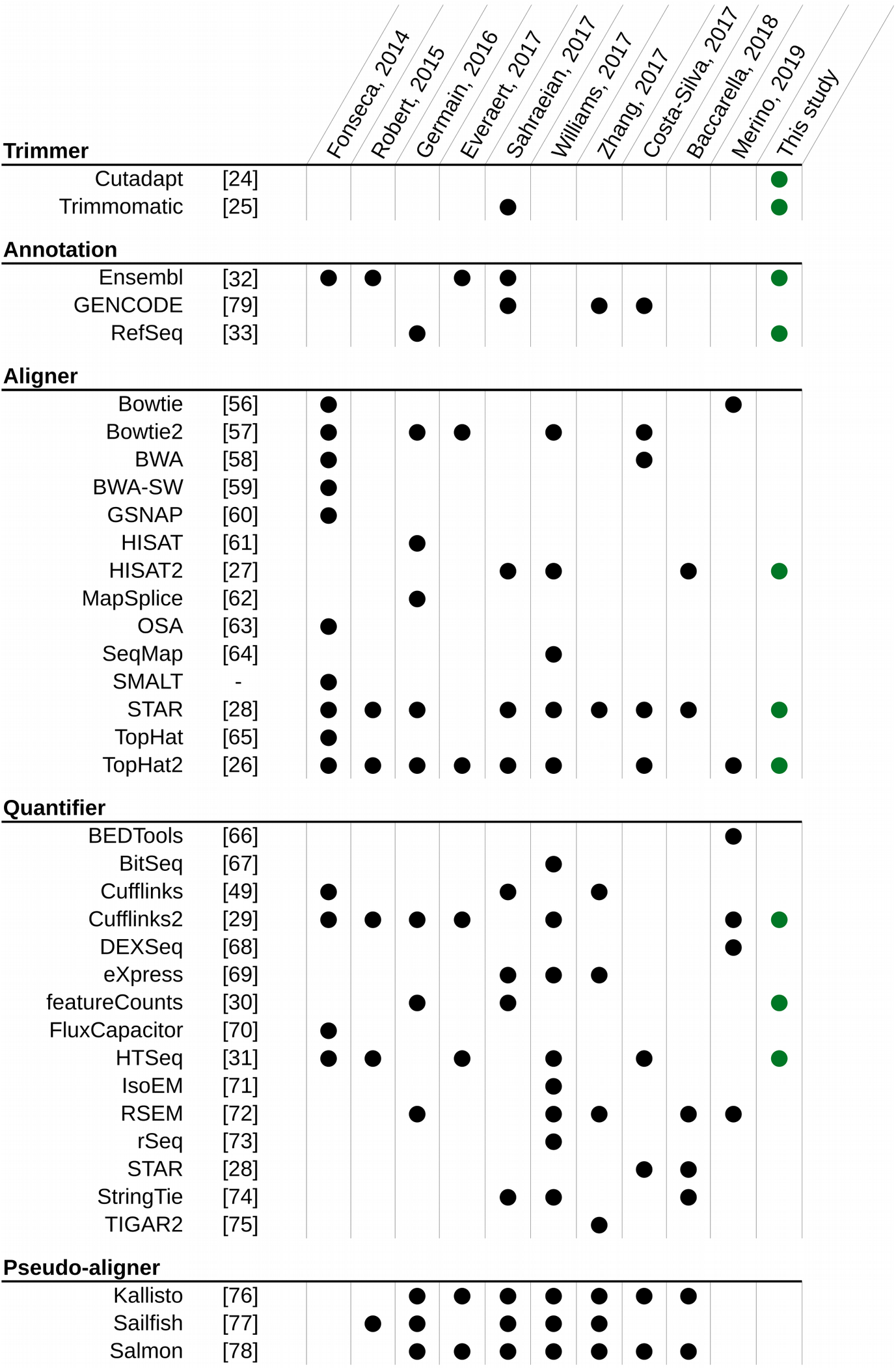
Software considered in RNA-seq workflow bencharking studies. Compilation of software and genome annotations used in articles benchmarking at least two different steps of the RNA-seq *in silico* pipeline. Software is classified by steps, where pseudo-aligners are considered separately because they overlap more than one step. Major re-release of software with an independent publication is considered as a separate software. The last column describe softwares considered in the present study.

Considering the difficulty in obtaining a high-quality gold standard for all genes in an RNA-seq study, we propose another strategy to identify biases in the processing pipeline. Instead of assessing the divergence in relation to the ground truth, we suggest treating this as a classification problem. If we process datasets with a variety of different pipelines, and then find some gene signatures classifying the processed datasets in accordance to some pipeline choice, then we would have identified processing biases affecting the quantification of genes.

Matrix factorisation methods are important tools for data-driven analysis, used to identify the main characteristics of highly dimensional datasets. Principal Component Analysis (PCA) is usually the go-to method used when confronted with such problems. PCA deterministically explains the variance of a dataset by projecting it onto decreasingly important latent variables, each constrained to be orthogonal to one another, uncorrelated. However, we chose to apply another method in this article, namely Independent Component Analysis (ICA). ICA decomposes a dataset into a specified number of latent variables of unconstrained variance, while optimizing for their independence, i.e. minimizing their mutual information. Both techniques produce uncorrelated latent variables, but only ICA produces independent latent variables when the original signal is non-Gaussian^18^. Where PCA highlights the largest trends present within the dataset, ICA seeks to extract independent structures, or phenomena, occurring within the dataset. This is due to a significant difference in the hypotheses at the core of these methods. PCA considers the data to follow a multivariate Gaussian distribution whereas ICA seeks a linear combination of non-Gaussian distributions. ICA has previously been applied to RNA-seq quantification results to infer groups of genes displaying a shared behaviour across several datasets^19,20^. ICA is a long-sought answer to the *cocktail party problem*, where an unknown number of persons talk in a room where a known number of microphones are placed^21^. The goal of the problem is to decorrelate the different microphone feeds to isolate the original speech of each person. The main hypothesis of an ICA is that every observation, microphone input, is a linear combination of a set of sources, herein the persons. In our case, the expression of genes acts as our microphone feeds, and each person can be thought of as a cellular process, a level of regulation, or a technical bias which has effect on the expression level of a subset of genes. By decomposing RNA-seq datasets generated using different treatments or biological origins, one can identify the sources, named expression modes in the context of RNA-seq studies^19^, that are important in defining and differentiating our datasets.

The biological importance of expression modes can be inferred by correlating them with known biological features. ICA has also been previously used to identify and remove batch effects by correlating expression modes with experimental features within the datasets^22^. We will dub the expression modes as either biological or technical modes, based on the types of variables with which they correlate. While technical modes correlating with sequencing batch might not hold information of interest for us, we hypothesize that this could be used as a tool to study biases in the RNA-seq *in silico* processing pipeline. By analyzing a number of datasets using a wide variety of different software and references, we could identify technical modes classifying the datasets by the pipeline used. These hypothetical technical modes would not appear in a normal RNA-seq experiment, where we do not usually use multiple software in parallel to accomplish the same step.

In this study, we processed biological replicates of different human tissues with a wide range of RNA-seq in silico pipelines obtained from the exhaustive combinations of selected software and genome annotations. We primarily chose software and references that are currently reported as being used in the RNA-seq literature, in order to represent the present situation^17^. We then decomposed these expression data into expression modes using an ICA analysis. We further characterized these modes as either biological or technical modes, on the basis of the variables that they can classify. Technical modes were then studied to explain the observed bias, identifying the features responsible for a different behavior of the software. A differential gene expression analysis was also produced for the different pipeline steps, highlighting the number of genes globally affected by these steps.

## METHODS

### Cartesian product of RNA-seq workflows

We used previously published RNA-seq datasets of human tissues from Array-Express E-MTAB-2836^23^ (https://www.ebi.ac.uk/arrayexpress/experiments/E-MTAB-2836/). Samples used are available in Supplementary Table 1. In order to have different levels of in-group and between-group variability, we chose four different tissues (colon, heart, testis, thyroid), each represented by four samples coming from different individuals. To evaluate the impact of every pipeline step, we processed all the datasets with a wide variety of RNA-seq workflows. We chose either recent, or commonly used, software and genomic annotations for each of the different RNA-seq steps considered, as defined in Figure 1. To keep every step independent from one another, we chose to exclude any software that encompasses more than one step (i.e. pseudo-aligners that overlap the alignment and quantification steps). We processed each dataset with the full compendium of Cartesian products of pipeline choices, meaning every possible unique combination of software and references. Using a design of experiments (DOE) terminology, this represents a full factorial experiment. To keep a basis of comparison, we used gene level counts as the output of the different pipelines. We only used tools which directly report gene counts, so there is no transformation of the final output.

**Figure 1.**
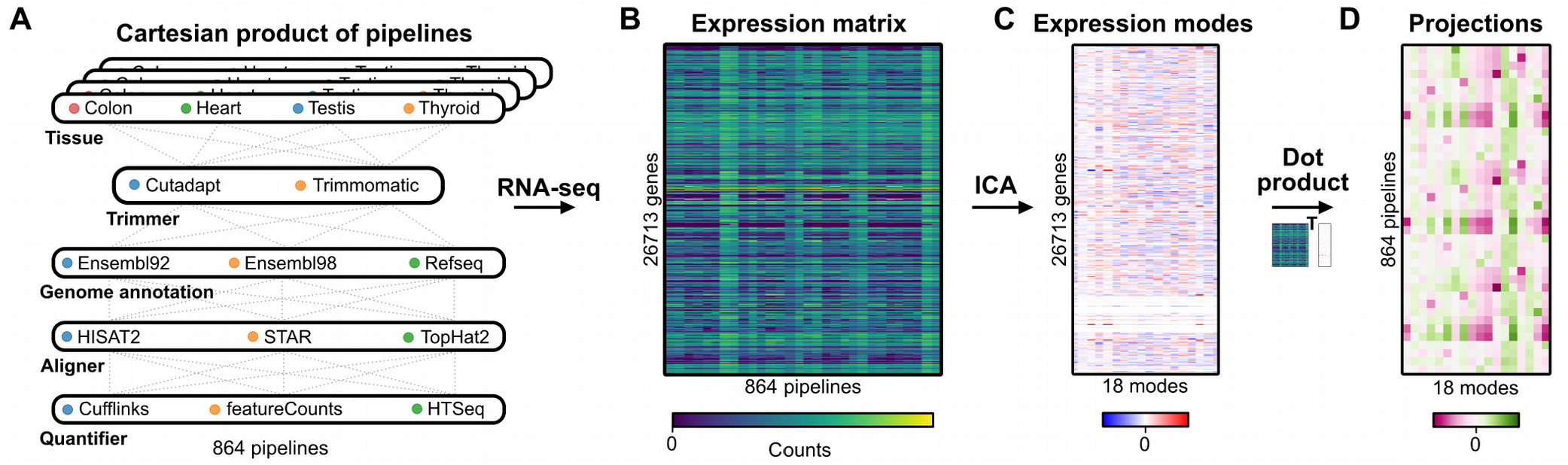
RNA-seq cartesian product and study design. Illustrations of the main steps of this study. First, all possible combinations of tissue samples, trimmers, genome annotations, aligners and quantifiers are processed as independent RNA-seq experiments. These results are compiled in an expression matrix which is decomposed into expression modes using an ICA. Projections, used to identify the information contained in the expression modes, is generated as the dot product of the expression matrix and the expression mode matrix.

### RNA-seq methodology

We performed RNA-seq using only methods that rely on genome-based alignment, and software that are restricted to a single methodological step. We kept default parameters for most of the options, as to mimic what is being done in the literature^17^. FASTQ files were downloaded from the SRA repository and trimmed independently using Cutadapt^24^ v2.3 and Trimmomatic^25^ v0.36. For the trimming parameters, we used a minimal Phred quality score of 15, and kept only reads that were at least 75 nucleotides after trimming, knowing that we are working with unstranded paired-end reads of 100 nucleotides. The alignment was performed independently using TopHat2^26^ v2.1.1 (wrapping Bowtie2 v.2.3.4.3), HISAT2^27^ v2.1.0 and STAR^28^ v2.5.3a. The aligners were run with default settings for unstranded data. It is important to note that only STAR requires an annotation file at this point. The other two software were not provided an annotation file for the alignment. The quantification was performed independently using Cufflinks^29^ v2.2.1, featureCounts^30^ (Subread v1.6.4) and HTSeq^31^ v.0.11.2. Quantification was summarized as gene-level counts. Ensembl^32^ version 92 and 98, and RefSeq^33^ release 109 were used as the different genome annotations. Ensembl 92 and RefSeq 109 were both released in April 2018, making them temporally comparable. Ensembl 92 and Refseq 109 are both built upon the GRCh38.p12 genome, while Ensembl 98 uses GRCh38.p13 genome. Because the primary assembly of both these reference genome versions is the same, and because we restricted our studied genes to the primary assembly, only GRCh38.p13 was used. Every time that a genome annotation was needed in a step, this step was processed three times, one with each annotation. The detection of differentially expressed genes (DEGs) was performed with DESeq2^34^ v1.26. All software tools were installed locally through Bioconda^35^. The dependencies and parameters for each pipeline steps are accessible in a Snakemake^36^ project.

### Data preprocessing

Raw counts from the different pipelines were combined in one expression matrix. Due to the fact that we are using more than one genome annotation, we require a common identifier to compare genes from Ensembl and RefSeq. To do so, we used the HUGO Gene Nomenclature Committee (HGNC) resource^37^. We considered data in the HGNC resource that were provided by Ensembl and the NCBI, while prioritizing information for HGNC in case of conflict. This also means that all results presented in this work are only drawn upon genes that are present in HGNC, ignoring genes that are unique to a specific genome annotation. After filtering for genes present in HGNC and quantified through the several pipelines, we are left with 26 713 genes. Raw counts were also preprocessed before being fed into the ICA model. For the first preprocessing step, we used the varianceStabilizingTransformation (fitType=‘local’) function from the DESeq2 project^38^. This step scales the different experiments so that they all have the same weight, and ensures the homoskedasticity of the genes, meaning that the variance of the genes is not function of their expression level. Homoskedasticity is important because the biological importance of a gene is not directly linked to its absolute expression value, and without this correction, the dataset features would be largely driven by a small number of highly expressed genes. The expression matrix is then transformed by a Mahalanobis whitening, rotating the dataset to decorrelate the different dimensions^39^.

### ICA model

We used the scikit-learn implementation of FastICA to process our dataset^40^. FastICA maximizes the neg-entropy, a measure of the non-Gaussianity of the components^21^. This optimization is performed using an iterative method, requiring the user to specify a tolerance, i.e. the minimum change of neg-entropy needed to stop the iterations. Because we need to perform FastICA with different numbers of components and due to the fact that the neg-entropy measure scales with the number of components, choosing a sensible tolerance is not trivial. A tolerance that is too large would end the optimization early, without attaining the real maximum, while a tolerance too small would never end the optimization. In order to avoid obtaining spurious maxima, we choose to force the FastICA algorithm to work with a really small tolerance (iteration step tolerance of 1e-18), and a large number of maximum iterations (1e5 iterations). While preventing the algorithm from stabilizing, we ensure that the optimization does not stop prematurely.

### ICA stability and independence

We then confirmed the robustness of the optimization maximum and the independence of the components. To do so, we ran the FastICA multiple times (n=25), using different starting points for the optimization. Afterwards, we computed the correlation matrix for the different components. In theory, an optimal correlation matrix for this problem should be a block identity matrix, where each block is a square with the same length as the number of replicates. Figure 2A contains an example of a near optimal matrix (with M=18), and two inadequate matrices (M11 and M26). The uniformity of the blocks confirms that the FastICA has found the same maximum for the different runs, and the identity matrix, where off-diagonal elements are zero, confirms the independence of the components. We scored the correlation matrix by quantifying its divergence from the optimal correlation matrix using the mean squared error (MSE).

**Figure 2.**
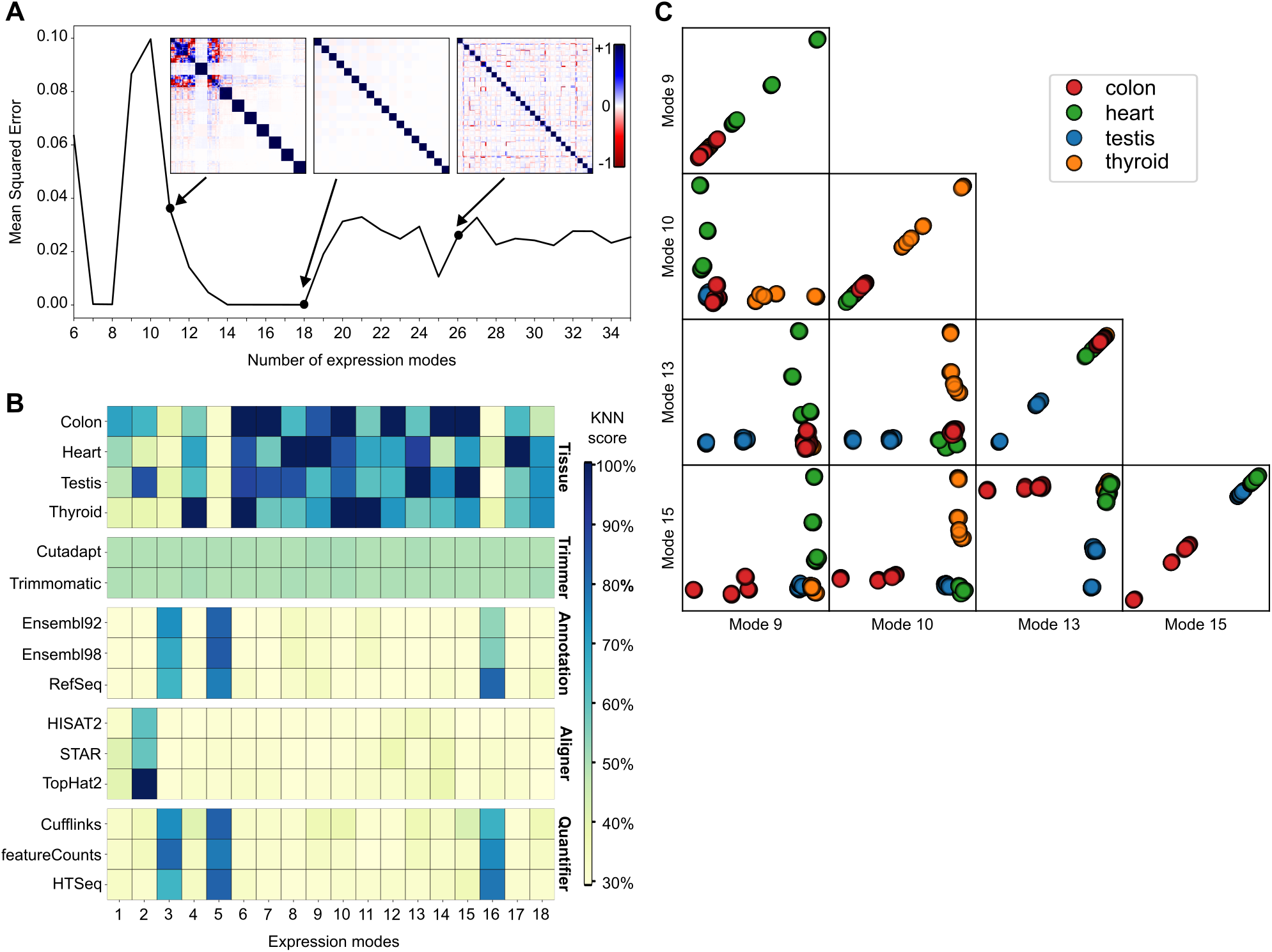
ICA decomposition of the RNA-seq data into expression modes. General information about the processed ICA model. **A** represents the Mean Square Error (MSE) for ICA model computed with different numbers of expression modes (M). The MSE is calculated using the theoretical optimal block diagonal matrix. Covariation matrix for models with M of 11, 18 and 26 are shown. **B** illustrates the information given by each expression mode. The heatmap is separated into five different blocks, each representing one variable choice. A high KNN score means that the variable is well clustered and well separated from the other variables. **C** illustrates distribution and pairwise distribution for four biological modes, one for each tissue type.

### Identifying expression modes

To identify what information an expression mode is providing, we used a k-nearest neighbour (KNN) classification approach. We first needed to generate projections of the pipelines along the expression modes. As in Figure 1, the projections are calculated as the dot product of the expression matrix by the expression modes matrix. This provided us with a one-dimensional distribution of the pipelines along each expression mode. We then quantified whether the pipelines are clustered according to biological or technical variables. To do so, using one projection at a time, and for each pipeline, we quantify the percentage of the 50 nearest pipelines, in terms of distance along the expression mode projection, that share the same label as the pipeline of interest for a specific biological or technical variable. This was done for all the different methodological choices from the different pipeline variables, taking the average percentage for each choice. This score informs us about the uniformity of the clusters found in a projection.

Each expression mode is defined by attributing a weight to each gene (Figure 1C), where genes with extreme values contribute more to the definition of the mode. In order to work with a list of genes, we needed to find a weight threshold at which genes would be considered as a part of the expression mode. We selected genes that were farther than four standard deviations from the distribution average, which creates gene groups with approximately 30 to 300 genes. The selected genes and their weights for all expression modes are available in Supplementary Data 1 for the original model and Supplementary Data 2 for the Cufflinks-only model. Only weights outside the four standard deviations were kept, the remainder were transformed to a zero value.

## RESULTS

### ICA highlights biological and technical differences between the RNA-seq pipelines

To run an ICA decomposition, one has to specify a number of expression modes (M) to be generated. This number is an unknown parameter and varies according to the underlying structure of the dataset. In order to identify the optimal number of expression modes to represent our dataset, we performed the ICA with a wide range for M (6 to 35), and we quantified the stability and the independence of the expression modes for these models. Figure 2A illustrates the distribution of the mean squared error (MSE) over the different number of expression modes used. Several values of M seem to be suitable for analysis, with a MSE of approximately zero. We chose to analyze the model with M = 18, being the model with the largest number of expression modes, while having the smallest MSE found. We favoured the largest number of stable expression modes with the hypothesis that decomposing the same dataset into more components would mean that the resulting components would be simpler, less convoluted.

We then identified the information given by each expression mode using the KNN score, displayed in Figure 2B. The heatmap is separated in five different blocks, each representing a pipeline variable, with the different choices, included in this study. For each block, the minimum possible score is 100% divided by the number of elements in that block, which is a score equivalent to random guessing. The heatmap should be read in a column-wise manner, looking at what information each expression mode is providing. The majority of the expression modes seem to be driven by biological information, which are the modes that are usually studied when using ICA with RNA-seq data^20,41^ and the modes of interest for researchers using RNA-seq to gain insight into biology. To confirm the informational value of the biological modes, we illustrated the distribution of the pipelines along four selected biological modes, one for each tissue, in Figure 2C. We can observe that each distribution offers a clear separation for a specific tissue, meaning that the underlying gene weights can be used as a biological gene signature for tissue classification. The pairwise comparisons also confirm that the modes are independent from one another because the pipelines are distributed as two perpendicular lines. We further characterized these gene groups using a Gene Ontology enrichment analysis provided in Supplementary Figure 1. These figures show that the genes used to classify the different tissues are also biologically related to the tissues, meaning that we have learned from biologically relevant features of the dataset. While the first block of Figure 2B represents the general use of ICA, i.e. studying biological features of a dataset, and serves as a positive control for our approach, our interest lies in the four other blocks.

The choice of trimming software, studied here using Cutadapt and Trimmomatic with the same set of parameters, does not have any impact on the dataset that could be identified by the ICA. The trimming block in Figure 2B shows a uniform block of value 50% for every expression mode, meaning that the distribution of the pipeline along the expression modes is purely random. While this does not prove that the trimming software does not have any impact, it shows that this impact would be smaller than, and therefore hidden by, the other expression modes.

The choice of alignment software is captured in the expression mode 2 (EM2), where TopHat2 is shown to be fully separated from the other software, whereas HISAT2 and STAR seem to be indissociable from one another, due to their similar scores. Genome annotations and quantification software are interlinked in three different technical modes (EM3, EM5 and EM16). Detecting technical modes is only the first part of the problem. Having shown that an ICA decomposition can be used to extract gene groups that seem to be specifically differently reported by different software, we next investigated whether a common feature in these gene groups can explain the differences.

### Discordant alignment of reads on gene-pseudogene pairs

Expression mode 2 (EM2) is composed of genes that differ in quantification in regard to the alignment software used in the analysis. Based on the KNN score of Figure 2B, these genes are similarly quantified when using either HISAT2 or STAR, but differently when using TopHat2. In Figure 3A, we can observe the EM2 weight distribution for all genes, where the genes considered significant (more than 4 standard deviations from the mean) are colored in blue and red, for positive and negative weights, respectively. In Supplementary Figure 2, we can observe the distribution of the pipelines along EM2. Seeing that HISAT2 and STAR have bigger projection scores than TopHat2, we can infer that the genes with a positive weight are more highly expressed when using STAR and HISAT2 than when using TopHat2, and vice versa.

**Figure 3.**
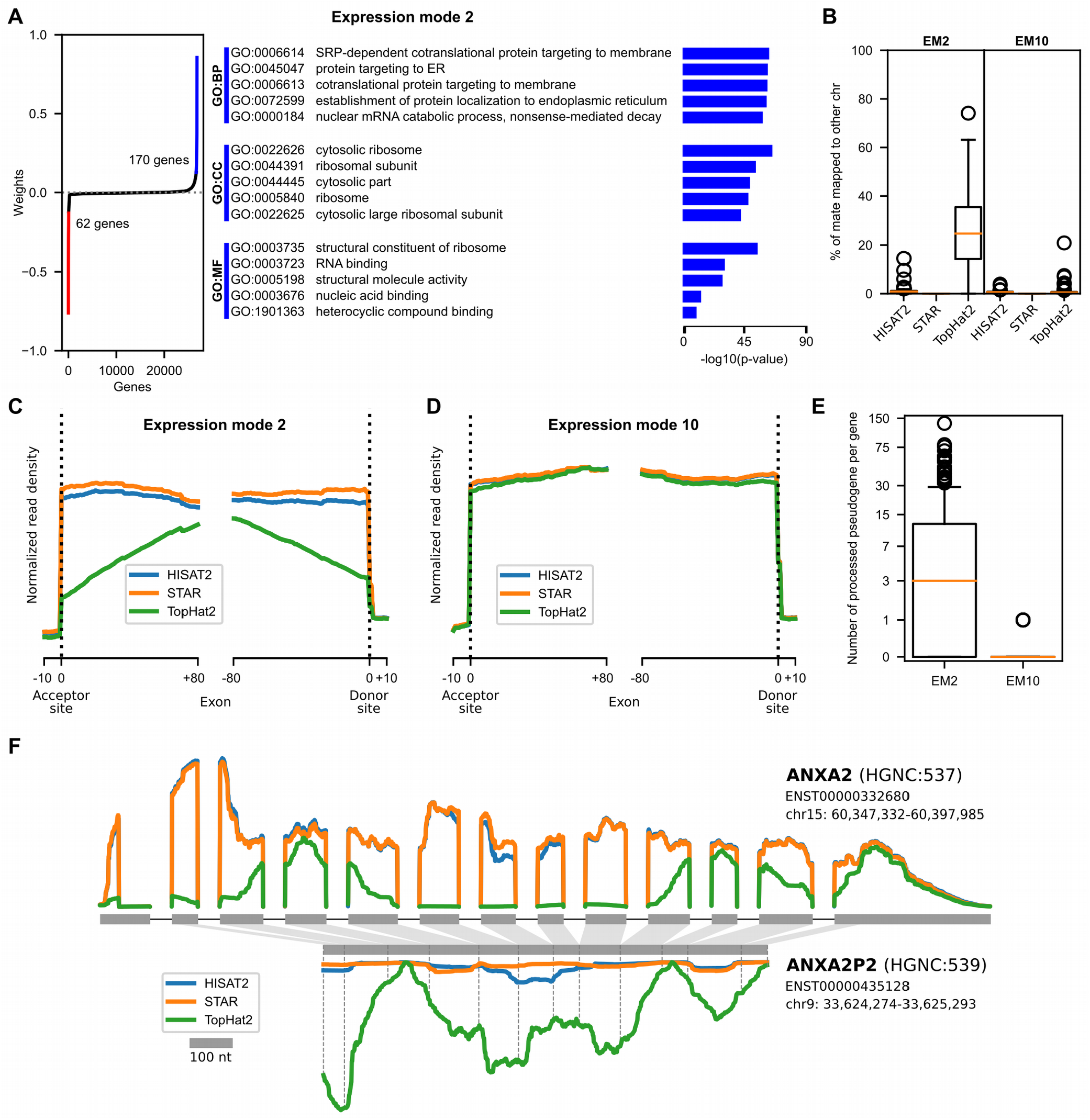
Technical mode linked with alignment software. Description of the features explaining the alignment software classification. **A** illustrates the gene weights distribution of EM2, along with their Gene Ontology (GO) enrichment analysis. Genes with a weight that is at least four standard deviations from the mean were considered significant and were coloured in the distribution. The GO enrichment analysis was performed using only the significant genes, and no enrichment was found in the negative genes. **B** shows the percentage of reads with mates that have been aligned onto another chromosome for the significant genes in EM2 and EM10, which is the negative control. The circles are the outliers of the boxplot. **C** is a metagene plot of the acceptor and donor sites of exon-exon junctions in EM2, with the profiles being separated by alignment software and **D** is generated using the EM10 genes. **E** quantifies the number of processed pseudogenes originating from the significant genes in EM2 in comparison to those in EM10. The plot is scaled using the inverse hyperbolic sine transformation. **F** describes the read profiles for each aligner along ANXA2 and ANXA2P2, the latter being the pseudogene of the former. The profiles are the averaged profiles of each considered pipeline and tissue combination. The exons are scaled accordingly to the 100 nucleotides (nt) reference in the legend. The introns were all truncated to a fix length in order to enhance readability. The mapping of the exons onto the pseudogenes was done using local sequence alignment, and position of the original splicing junctions are marked with dashed lines on the pseudogene. The pseudogene read profile is presented upside-down, and the two read profile plots are presented using the same scale.

Figure 3A also displays results from a Gene Ontology (GO) enrichment analysis of the significant genes. This analysis was done separately for the genes with positive and negative weights. Only results of the enrichment for the positive genes are shown since the analysis of the negative genes led to no significant enrichment. Interestingly, we found a strong enrichment for ribosome and translation related GO terms in the positive genes group. For this to happen, the common feature of the positive genes that is considered differently by the alignment software must also be linked to some biological characteristics of the genes. By exploring the average alignment statistics for the positive genes in EM2, we found that TopHat2 has a significant percentage of mapped read pairs with a mate aligned to another chromosome, as displayed in Figure 3B. For the same genes, STAR reports no read pairs mapped to different chromosomes, and HISAT2 has an average of 1% (in comparison to 25% for TopHat2) of mapped read pairs in this situation. In order to identify whether the observed effect is expression mode dependent, we used another unrelated expression mode as a control. By comparing these results to genes from a biological thyroid-related expression mode (EM10), we do not find the same effect. In EM10, the three software tools have a similar and a nearly null number of read pairs mapped to different chromosomes. Having identified a divergent characteristic, we then analyzed the discordant read pairs, by comparing alignments from STAR and HISAT2 to alignments from TopHat2.

Metagene plots in Figure 3C and 3D illustrate the aggregation of read profiles for all exon acceptor and donor sites for the gene groups of interests. While Figure 3D shows a very similar read profile for all alignment software, Figure 3C shows a dissimilar profile for TopHat2. Thus, HISAT2 and STAR profiles are similar in EM2 and EM10 but the TopHat2 profile in EM2 lacks reads around the exon-exon junctions. A theoretical exon-exon junction profile would show a perfectly square profile at the acceptor and donor sites. In our case, the progressively smaller profile, as we approach the edge of the exon, is a sign of a difficulty to align reads spanning across an exon-exon junction. Because we are aligning on the genome, alignment software must be able to map reads in a discontinuous manner across an exon-exon junction, namely gapped alignment. All three tested aligners are known to perform gapped alignment, but TopHat2 seems to fail to do so in the specific situation highlighted by the ICA expression mode. The biological particularity of the EM2 positive genes is that they possess a significantly higher number of processed pseudogenes in comparison to the other genes, as illustrated by Figure 3E. The gene-pseudogene relationship used is described by the PsiCube project^42^. Processed pseudogenes are defined as the product of the retrotranscription of spliced RNA inserted back into the genome. This means that they have the genomic sequence of the transcribed product of their parent gene, i.e. continuous exon-exon junction sequence. In our situation, TopHat2 prefers using a distant already spliced junction than using a local junction that needs splicing. The creation of processed pseudogenes has been shown to favor highly conserved genes that are widely expressed^43^, without any detectable sequence bias^44^. This definition fits well with the ribosomal proteins and translation associated machinery found in the GO enrichment analysis.

### Expression modes each identify gene groups with opposite behaviours

Having demonstrated that the positive genes in EM2 have lower read mapping in TopHat2 due to the presence of pseudogenes, we turned our attention back to EM2 negative genes. Due to the two-tailed distribution of gene weights, we would expect an opposite effect, meaning that the negative genes should have a higher read mapping in TopHat2 than HISAT2 and STAR. We should also point out the abnormal (when compared to the other weights distributions) shape of the negative gene weights, harboring a very steep change of weight, instead of the progressive asymptotic-like shape of the distribution. Using Ensembl 98 biotypes, we found that the majority of the negative genes are pseudogenes (44/62) and the remainder are protein-coding genes (18/62) and those are primarily mono-exonic (13/18). In Figure 3F, we illustrate a pair of genes, AXNA2 and AXNA2P2, that were both found in EM2, both in opposite gene groups. AXNA2P2, as its symbol indicates, is a pseudogene originating from AXNA2, and the correspondence between the two, established with local alignment of their mature RNA sequences, shows that the pseudogene is a truncated intron-less copy of a transcript from the original gene. Averaged read profiles from all considered RNA-seq pipelines, separated by alignment software, are shown for both genes. TopHat2 profiles are obviously quite different from the other aligners, but more interestingly, both of its profiles seem to overlap, where the sum of the two profiles is similar to HISAT2 and STAR profiles of the principal gene. For example, the peak of TopHat2 reads on the ANXA2 fourth exon is related to the minimum value found in the ANXA2P2 corresponding section. This figure provides additional proof that TopHat2 shares the reads between a gene and its pseudogenes, while HISAT2 and STAR do not. Supplementary Figure 3 provides three other pairs of genes illustrating the same situation. To further our argument, we can observe that the exon length is responsible for part of the TopHat2 profile. In the RPL13A profile, the exons are too short for reads to be mapped exclusively onto one exon, creating a situation where the pseudogene is getting nearly all the reads. Conversely, GLUD1 has longer exons, and we can observe the same exon profile as found in the metagene plot Figure 3C. GLUD1 also shows, looking at its last exon, that as soon as we are in an exon long enough to map entire reads, the three aligner profiles converge. We believe that the steep change observed in the weight distribution originates from having a binary feature which is being a product of a spliced retrotranscription or not.

### Genome annotations and quantifiers interact in RNA-seq quantification

Expression modes 3, 5 and 16 offer a more convoluted story, because both the genome annotations and the quantification software have been partly clustered in them. This means that these two steps, for the genome annotations and software selected in this study, are not fully independent. In Supplementary Figure 4, we can observe the distributions of the different pipeline projections along the three technical modes of interest, colored by their annotation and quantification software. The same pattern can be seen in the three technical modes. In both steps, we can find two different, loosely defined but clearly separated, clusters. One cluster always contains two genome annotations and the same two quantifiers, featureCounts and HTSeq, while the other cluster contains the three genome annotations and the three quantifiers. Because quantification is downstream of the genome annotation specification, we will illustrate this phenomenon as being quantification driven. The hypothesis here is that there are some features in the genome annotations upon which a classification can be done if the pipelines used featureCounts or HTSeq. These different features can also be used to generate all three possible binary classification of annotations. However, when using Cufflinks as the quantification software, these features appear to have no effect on the quantification, because Cufflinks is shown in a single cluster. Therefore, Cufflinks seems to rescue the problematic features which cause quantification biases when using featureCounts or HTSeq.

### Expression modes may hide other gene groups with similar quantification power

This situation lets us test a hypothesis that we spelled out earlier, which is the fact that some expression modes can hide other, smaller, expression modes. To test this, and to also put Cufflinks to the test, we generated another ICA model, in which we only used expression datasets that were generated using Cufflinks. This ICA model was processed in the same way as previously described, with M = 16 being the model with the smallest MSE and the highest number of expression modes. The KNN score heatmap of this ICA model can be found as Supplementary Figure 5A. In this Cufflinks-only ICA model, we can find two different technical modes, independently related to the annotation and the aligner blocks. Both modes seem to have the same classification power (KNN score pattern over the different tools in a block) as another technical mode found in the original ICA model. To verify whether we have found the same expression modes in two different models, or expression modes based on different gene groups displaying the same classification power, we can compare the overlap of significant genes in both expression modes. This overlap is shown in Supplementary Figure 5B, where M2 is compared to MC7, both being similar alignment technical modes, and where M16 is compared to MC14, both being similar annotation technical modes and both able to separate Ensembl from RefSeq. In the first case, we can observe that the genes from the two alignment technical modes largely overlap, which means that they have probably learned from the same gene groups, using the same features. Conversely, the technical modes describing the annotations have a small overlap, meaning that the two modes have been built on mainly independent gene groups, which also means different features. The overlap might be explained by genes having features that let them be part of both groups. We therefore have two different expression modes, dependent on the quantification software used, that have classification power over RefSeq and Ensembl genome annotations.

### A gene quantification is affected by its definition and its neighbouring gene definitions

Figure 4 explains the main features used by expression modes 16 and C14 to drive the clustering. Expression mode 10, the thyroid-related biological mode, was also used as a negative control, because it is not expected to be enriched for the annotation classifying features. First, we looked at the extent of exon overlap for the genes considered, within each annotation. To do so, we measured the proportion of exonic nucleotides that overlap any other gene in the given annotation, considering both sense and antisense strands (outer most plots in Figure 4). These analyses indicate that negative and positive significant genes in M16 are exhibiting an opposite pattern which is not found in MC14 and M10. The negative M16 genes are highly overlapped in RefSeq while being marginally overlapped in Ensembl 92 and 98, and inversely for the positive genes. Next we compared the annotations pairwise and measured the percentage of unique and common sequences for the genes of interest when two annotations are compared (flanking inner plots). These analyses show that MC14 is now also exhibiting a mirroring pattern, where its negative significant genes are longer and have more unique sequence in Ensembl than RefSeq, while the opposite is true for its positive genes. M10 shows that some baseline of sequence differences might be expected, with Ensembl having more unique sequences than RefSeq, but only C14 has a clearly defined mirror effect. This brings us to the observation that, because the ICA generates twotailed distributions for the gene weights, we are able to see genes exhibiting both extremes of a feature, as clearly shown by the mirroring effect in Figure 4. While it is clear that both technical modes have a main feature, where M16 is mainly affected by overlapping annotations and MC14 by different gene definitions, we can also see aspects of the other feature in both sections. This might be due to the fact that the two technical modes share some of their genes, leading to genes contributing in both sections. One can also easily imagine how having a longer gene sequence might also result into having more overlapping sequences, showing that these two features are not completely independent.

**Figure 4.**
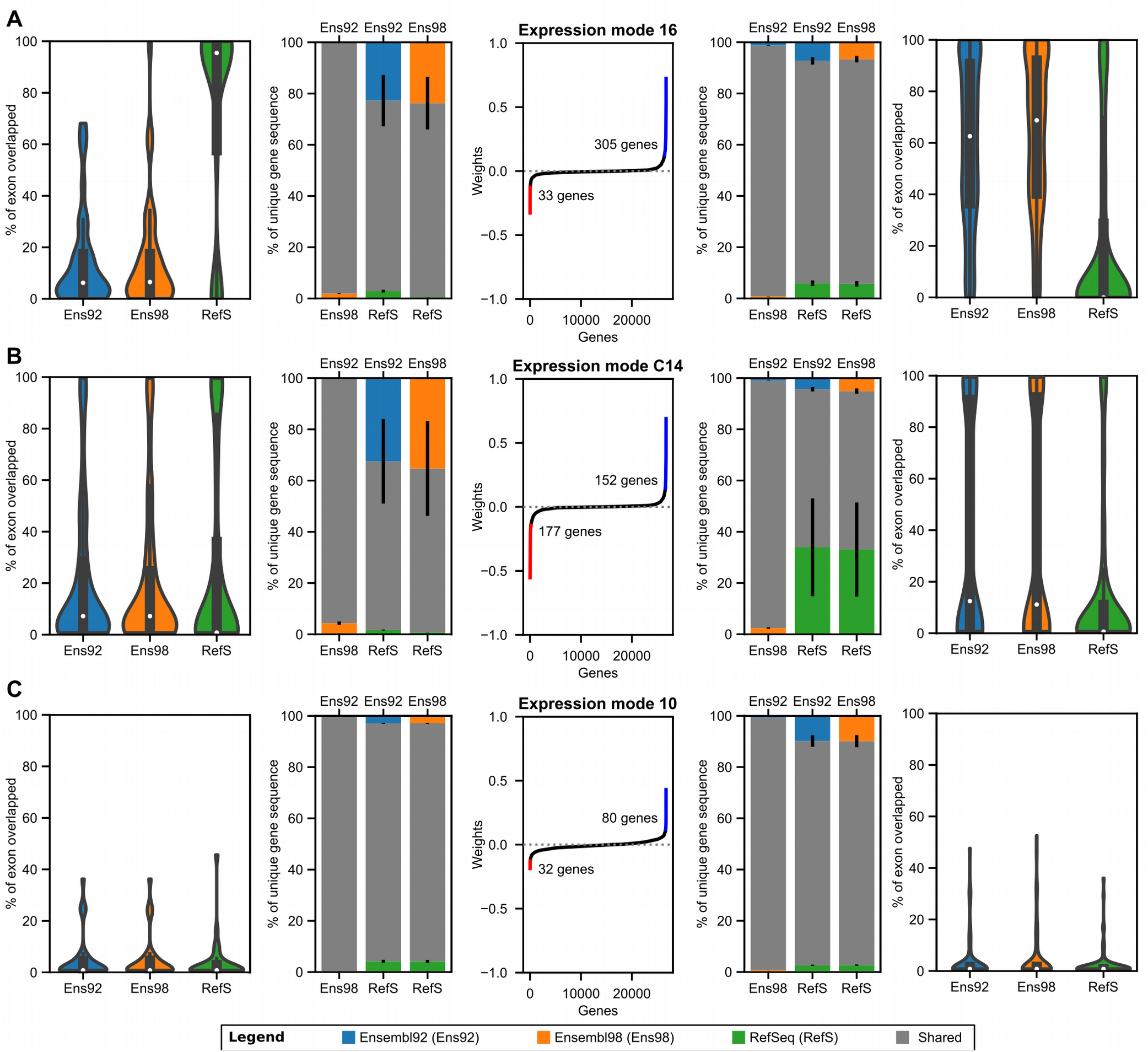
Technical modes linked with Ensembl versus RefSeq classification. Description of the features leading to an Ensembl versus RefSeq classification in our expression datasets. **A** and **B** are technical modes linked with genome annotation classification and **C** is a negative control, a biological mode linked with thyroid-related gene groups. The middle plots describe the distribution of gene weights in the mode, highlighting the genes considered to be significant (more than 4 standard deviations from the mean). On both sides of the middle plot, sets of similar plots are found. The plots on the left show metrics calculated from the significant genes with a negative score, and conversely for the right plots. The outer most plots show the distribution of the percentage of a gene exons, by nucleotide length, that are overlapped by exons from other genes. Overlapping genes from the sense and antisense strands were both considered. The inner flanking plots show the average of shared and unique exon sequences for each pairwise combination of genome annotations. Sections drawn in an annotation color represent the percentage of sequence not found in the other annotation, while the shared section can be found in both annotations. The black lines represent the error bars. For both flanking plots, each individual genomic position was only considered once, independently of the number of individual exons of the same gene that may overlap that position.

### Extrinsic and intrinsic factors of genome annotations affect quantification differently

Interestingly, the quantification software behaviour classifies them by the approach that they use to solve the quantification problem. Both featureCounts and HTSeq are count-based quantifiers, whereas Cufflinks quantifies through transcript assembly and quantification. We have observed that the main bias of count-based quantification is the definition of neighbouring genes, seen through the metric describing the percentage of exon overlap, whereas transcript quantification is primarily affected by the definition of the gene itself. We will describe the former bias as originating from the extrinsic factors, and the latter from the intrinsic factors. By projecting expression datasets from featureCounts and HTSeq onto the technical mode C14, as seen in Supplementary Figure 5C, we can see that these two tools are also being affected by the intrinsic factors, because they are shown to cluster in the same way as Cufflinks does. The fact that MC14 does not appear in the original ICA model means that M16, the extrinsic factors affecting featureCounts and HTSeq, exhibits a stronger bias that is hiding the bias caused by intrinsic factors. We can also pose that the intrinsic factors are independent, meaning that all quantification software behave the same way towards them, whereas the extrinsic factors are the ones responsible for the interaction between genome annotations and quantifiers, because they are treated differently by the software.

Supplementary Figure 6 illustrates three groups of example genes that were found to be significant in at least one of the two technical modes classifying Ensembl and RefSeq. All three genes of interest are protein-coding genes with conserved consensus coding sequence^45^ (CCDS) across all three genome annotations. The first gene, ARPC1A, was found in EM16 and is a good example of a gene being overlapped by a read-through gene that is unknown from RefSeq. The second example, GOLGA8M, was found in EMC14 and is shown to have major differences in annotation from Ensembl to RefSeq. There are also some overlapping genes that are not the same, but they are smaller in comparison to the overall gene. Intriguingly, Ensembl possesses two different genes (ENSG00000188626 and ENSG00000261480) that have the same gene symbol and reported HGNC ID, while HGNC only lists the first as being GOLGA8M. Moreover, the first has two transcripts which are named GOLGA8M-201 and GOLGA8M-202, while the second gene only has GOLGA8M-203. The last example, KCNA6, was found in both EM16 and EMC14 and exhibits both a change in overlap and a change in definition across Ensembl and RefSeq. This figure provides a better understanding of the separation between extrinsic and intrinsic factors.

### ICA can correctly classify different versions of the Ensembl genome annotation

We have shown that it is possible to classify RNA-seq quantification results with respect to their source of genome annotation. To delve deeper into this issue, we have also included in the study two different versions of the Ensembl genome annotation, versions 92 and 98, which were respectively published in April 2018 and September 2019, approximately 18 months apart. We wanted to see whether we could also classify them, and if applicable, identify the way they diverge from one another. The ICA model has identified two technical modes, EM3 and EM5, that differentiate between the two Ensembl versions, with RefSeq pairing with a different Ensembl version in each technical mode, as seen in Supplementary Figure 4. Once again, Cufflinks does not behave as the two other quantifiers, affecting the global clustering of the annotation projections. From our previous observations on EM16, we can hypothesize that EM3 and EM5 classifications will also be driven by extrinsic factors, meaning genes with overlapping loci. Supplementary Figure 7 presents the data supporting our explanations of the clustering, with the same plot representations as used with the previous technical modes. However, both technical modes do not offer the same prominent mirroring effect as observed in Figure 4. If these modes are also driven by extrinsic factors, it would be expected that the outer plots show a clear difference in the percentage of exon overlap. In EM5 (Supplementary Figure 7B), positive genes display a clear separation between Ensembl 98 and the other annotations when quantifying the percentage of exon overlap. Notably, the distribution of scores for the two other annotations is approximately the same, as it was for the same plots in Figure 4. With the mirroring hypothesis, we would expect that Ensembl 98 would exhibit a smaller percentage of overlapped exon in the negative genes, which it does. However, its distribution is not clearly different from Ensembl 92, and Ensembl 92 and RefSeq distributions do not look alike. Both of these points show a divergence from the previously observed data. In order to explain this discrepancy, we characterized the difference in percentage of exon overlap for each gene in the components, across the genome annotations. Supplementary Figure 7C shows this characterization for both EM3 and EM5, the plots being relative to the annotation that is clustered alone, which is respectively Ensembl 92 and Ensembl 98. Positive and negative genes are split into three groups, with respect to their difference in exon overlap with the reference annotation. If both scores of a gene are within 10% of the reference annotation score, the gene will be put in the middle group, being approximately the same as the reference annotation. If the gene has a score of at least 10% higher or lower in at least one annotation, it will be classified accordingly in the bigger than or smaller than group, while also being colored relative to which, or both, annotation diverges from the base case. If both scores are outside the 10% threshold in opposite directions, the gene will be classified as similar to the reference annotation. Looking at the results for EM5, we can see a big difference between the positive and negative genes. Based on the mirroring, it was expected that the genes would be separated mainly in a diagonal fashion, but the new information that we gain from this representation is the unequal separation of the exon overlap score across the two annotations. While the positive genes are mainly below the threshold for both annotations, the negative gene group is dominated by genes that are only differing in RefSeq. The same kind of grouping can be observed in EM3, where the majority of the genes contributing to the exon overlap score are only positive for one annotation. The positive genes plot displaying percentage of unique gene sequence in EM3 also shows that there might be some influence of intrinsic factors as well, with Ensembl 98 and RefSeq having more unique sequence compared to Ensembl 92. Finding large enough differences to enable classification of two different genome annotation versions is a much more difficult task than for two different annotations, and the technical modes demonstrate this by using different sets of genes to be able to cluster differently. We believe that these gene groups are heterogeneous groups, with each subpart contributing to differentiate with one annotation at a time.

### Ensembl distinguishes itself by a higher, and growing, number of overlapping loci

EM3 and EM5 let us explore the differences in Ensembl, and posit in the way that Ensembl is currently evolving through time. Interestingly, the clarity of the gene groups observed in Supplementary Figure 7C may be used to interpret the evolution of the annotations. Based on the projections, Ensembl 98 is less expressed in the positive genes, and to achieve that, we only need to find genes that have gained overlapping loci in Ensembl 98 and that are not overlapped in RefSeq. Conversely, to be more expressed in EM5 negative genes, Ensembl 98 needs to lose an overlapping loci from Ensembl 92, loci that must be present in RefSeq. From the results, we can conclude that it is easier for Ensembl 98 to gain new overlapping annotation than to lose some from Ensembl 92, and we can also conclude that RefSeq seems to globally possess fewer overlapping genes, because it is mainly responsible for the EM5 negative genes. In order to explore this, we quantified the percentage of overlapped exon across all of the 26 713 genes considered in this study. This quantification is shown in Supplementary Figure 7D, as distributions only including genes that have a non-null overlap. Quite surprisingly, the distributions for the different genome annotations are essentially that same, made from a different number of genes. While anecdotally looking at the genes responsible for the new overlaps in Ensembl 98, we stumbled upon many read-through, that were not necessarily identified as such, and set forth to quantify them in Supplementary Figure 7E. We defined a read-through as a gene having at least a transcript that has, for a least two different genes, a perfectly matching exon, based on the genomic coordinates, and not the sequence alone. These matching exons must also be distinct exons in the overlapping transcript. We also only quantified read-throughs that are overlapping at least one gene included in our study. When comparing Ensembl 92 and RefSeq, based on the last two plots, we can acknowledge that Ensembl has far more genes with overlapped loci (38% against 27% for RefSeq), and about three times more read-throughs than RefSeq. We can also see that Ensembl 98 is straying further away from RefSeq, with Ensembl 98 having even more overlapped genes and read-through genes.

### Differential expression analyses describe the same extent of methodological step biases

A differential expression analysis (DEA) is usually used to identify whether a gene expression is varying accordingly to an experimental variable^46^. In comparison to the ICA analysis that we performed, DEA reports all genes independently, whereas gene groups in the ICA have some shared expression patterns. In order to compare the genes found through the ICA to a more commonly used technique in the RNA-seq field, we performed several DEA using the pipeline variables and their choices as respectively experiments and conditions. For example, to generate a DEA on the impact of the trimming, we compared expression datasets processed using Trimmomatic to those processed using Cutadapt, using all the corresponding datasets as replicates. This means that our replicates are very heterogeneous, having expression datasets grouped together that were generated using the whole spectrum of the other pipeline steps. In a DEA, negative results do not mean that the genes were not impacted by the condition, but that such an impact could not be observed within the variance of the datasets. Because some pipelines do have a significant impact on the quantification, other steps might suffer from large within-group variance, which makes them less likely to have significant results. Our multiple DEA analyses have very heterogeneous within-group variance due to the fact that we are reanalyzing the same expression datasets, with different groupings of the samples with respect to the different methodological steps.

The number of replicates is usually a variable worth optimizing in DEA, since more replicates means more sequencing, and sequencing is still a costly task^47^. In our case, the replicates are mainly generated through processing the same dataset using a different *in silico* pipeline, meaning that we have an abnormally large number of replicates for the different experiments. These replicates translate into abnormally small p-value, and we even hit the number limit of a 64-bit system, where number smaller than approximately 10e-308 are considered as 0 and reported by DESeq2 as such^34^. In the volcano plots, any gene with a p-value of 0 was displayed as having (in −log10 form) the maximum possible value.

Figure 5 displays volcano plots for the different technical DEA, grouped by pipeline step. Using a p-value of 10e-35 and a log2(fold change) of 2 as our significance thresholds, we have identified the significantly differently expressed genes (DEGs) for the different experimental choices using their colors and displayed their counts on the lower edge of the volcano plots. It is apparent from an overview of the different pipeline steps that the methodological choices do not bear the same importance in the definition of the results. We can also immediately see some resemblance with the ICA results. The trimming step is the only step not highlighted by the ICA, and all of its genes in the volcano plot are centered around the null coordinates, nowhere near significance. Accordingly, the alignment step was captured by only one technical mode in the ICA model, and has the second-lowest number of DEGs. It can also be observed that HISAT2 and STAR generate results that are more similar than TopHat2. And lastly, genome annotations and quantifiers seem to have a similar number of DEGs, and the same tool clustering as found in the ICA is observable.

**Figure 5.**
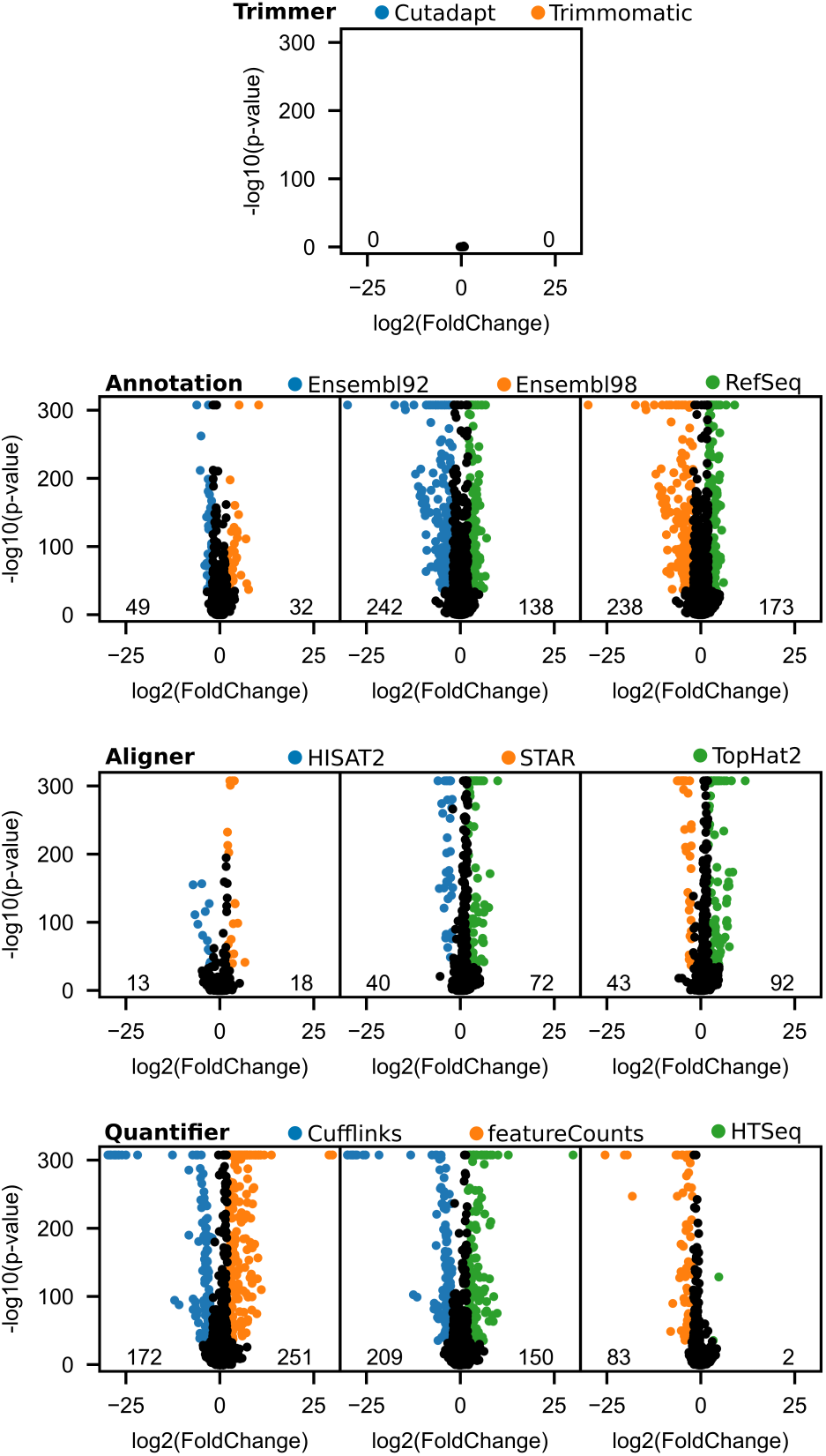
Differential expression analysis of RNA-seq methodological choices. A differential expression analysis was performed for each pairwise combination of choices for each pipeline step. The volcano plots of these analyses are presented here, where significantly differentially expressed genes (p-value < 10e-35 and log2(fold change) >= 2) are colored accordingly to the choice in which it is overexpressed. Numbers on the lower edge of the plots quantify the number of colored genes.

## DISCUSSION

To our knowledge, the community has overlooked the “what-to-benchmark” question regarding the RNA-seq quantification pipeline^48^. This question might be more akin to a manufacturing context, where optimization of resources is directly linked to the metric of success, whereas it can be more fuzzily linked in research. The “what-to-benchmark” question is in fact of utmost importance in any resource-limited context, such as research, to correctly identify the most pressing questions. In the context of RNA-seq, the “what-to-benchmark” question is related to the action of benchmarking the sensitivity of the pipeline as a whole, in order to identify the greatest source of variability in the results.

In this study, we described quantification biases found in the whole RNA-seq *in silico* workflow, using an ICA decomposition on expression datasets created through reprocessing of RNA-seq experiments by a wide variety of methodological choices. We put forward the idea of technical reprocessing to identify biases between different choices of a specific methodological step, instead of assessing the deviation from the ground truth, which is a difficult value to find. Biases in a specific methodological step are proof that developers either used different biological hypotheses or have encountered unknowns. While this approach is not a panacea, since it will fail to identify any problematic shared by all the methodological choices, it can be an interesting technique to characterize the current landscape of quantification discrepancies concerning tools presently used in the literature. Such landscape can be characterized by a within step manner, which is the usual analytical benchmarking approach, but also a between step manner, which highlights the relative importance of the different methodological choices. The idea of gene specific biases identification in a methodological context in not new, but we extent this approach by being able to link similarly affected genes together, leading to an easier identification of impactful characteristics of problematic genes^7^.

While the format of this study does not let us provide direct recommendations regarding what methodological choices to prefer, we can highlight choices that we would not recommend. While Tophat2 is still one of the primary aligner software in use in the literature^17^, we cannot recommend its usage due to its bias towards genes with processed pseudogenes. Furthermore, HISAT2 is a reimplementation of TopHat2, correcting some issues that have become apparent through the years. Authors of TopHat2 have also published many times, as seen in TopHat2 and HISAT manuals, and on the twittersphere, about the need to move towards newer software. While further models might highlight discrepancies between STAR and HISAT2, the data shown here cannot point towards one or the other.

We cannot recommend using count-based quantifiers, such as featureCounts and HTSeq, since, as stated by themselves^30^, they are not able to give accurate estimation of isoform quantification. The issue is that the isoform quantification problem is actually an overlapping transcripts problem, which means that it is not limited to transcripts of a single gene. If no genes were overlapping, every read falling into the region of a certain gene could be trivially assigned to it. But, as demonstrated by the technical modes linked to the quantification software, gene-level quantification for count-based quantifiers fails when the genes are overlapped by other genes. This also means that count-based quantifiers have overlooked the fact that genes may share some genomic coordinates. Based on that information, it would make sense to recommend using quantifiers that quantify on the transcript level through techniques trying to infer the transcripts from which each read could have been produced, a class of quantifiers which is represented by Cufflinks in this study^49^. As we have demonstrated, such software seems to be less affected by genome annotation extrinsic factors.

As for the choice of genome annotations, the recommendations are not as clear. Genome annotations do not hold the same place as the other software tools in the RNA-seq pipeline. Performance of software can be tested under specific conditions, and divergence from the expected behavior can be assessed. This was highlighted by technical modes, for example the lack of reads on exon-exon junctions for some genes when using TopHat2 contradicts our understanding of gene expression and the hypothesis of uniform distribution of reads along a gene. But genome annotations are information resources, having the dual purpose of being a repository of our gene biology knowledge, and a research tool, leveraging high-throughput techniques. While the technical modes associated with genome annotations informed us of differences in the annotation of different genes, differences that have a variety of impact on the basis of the quantification software used, it is difficult to assess which genome annotation is closer to the truth. Learning about differences that are driving the main biases in quantification is important for our understanding of the place of genome annotations in the RNA-seq pipeline, and while it does not give a clear answer about which annotation to use, it informs users about the non-triviality of choosing an annotation, and about features that are important to look for. On the other side, it is not that the annotations have converged on a similar understanding of the structure of a gene that this convergence is in phase with our current understanding of biology. To illustrate this, we can take the extreme example of the GDF1 and CERS1 genes that individually have an identical gene-level structure across Ensembl92, Ensembl98 and RefSeq, while also sharing the vast majority of their exons, as seen in Supplementary Figure 8. Overlapping loci are difficult to quantify, which makes these genes susceptible to unreliable quantification across the methodological landscape, but it also raises the question of whether these two entities are really independent genes. GDF1 and CERS1 even share an identical CDS in Ensembl through two transcripts (GDF1-201 and CERS1-205) that only differs by some nucleotides in both UTR extremities. It was proposed that GDF1 produces a polycistronic mRNA^50^, but the two proteins discussed in the paper are now products of the two different genes. Genome annotations are not a data structure that is currently able to support polycistronic RNA, and it might be what justified the separation into two different genes. But knowing that overlapping genes cause issues in quantification, and that, if they are truly polycistronic, there is only one RNA to quantify, we should review the genome annotation information structure. There have been multiple projects exploring the polycistronic nature of human mRNAs (e.g.^51,52^). The inclusion of such data would require allowing transcripts to possess multiple CDS elements. This is simply an example that biological understanding and hypotheses evolve, and that our software, and usage of them, must follow accordingly.

To compare the genome annotations, we had to limit ourselves to genes that have some basis of comparison, and HGNC was used to bridge the gene identification between RefSeq and Ensembl. Since not all genes are annotated into HGNC, and since not all HGNC genes have an identifier for both genome annotations, our number of comparable genes is smaller than the total number of genes, and one could hypothesize that the remaining genes are better described and are probably more similar than the other genes. Furthermore, some of the remaining genes were not reported by the pipelines when using RefSeq. These genes, the 37 mitochondrial genes and 10174 pseudogenes (list available as Supplementary Data 3), are present in the RefSeq GTF annotation file, but are described using a single ‘gene’ feature, instead of the expected hierarchy where each gene possesses one or more transcript, themselves having one or more exon. In the RefSeq GFF3 file, some of these genes appear to have the expected hierarchy, but they do not respect the same naming scheme as the other genes (NM and XM identifiers for transcripts), and they are always mono-exonic. This issue has many implications. First, one would expect to have the exact same information within the different file formats distributed by a centralized resource. Second, deviating from the expected data format can create unexpected behavior for data processing software, seen here as genes not being quantified. Third, pseudogene is a very wide RNA category which includes mono-exonic processed pseudogenes, but also intron-bearing unprocessed pseudogenes. This makes for an unfair comparison if we were to compare quantifications of unprocessed pseudogenes between Ensembl and RefSeq. Supplementary Figure 9 provides such a visual comparison for the difference in gene structure. But because we do not have RefSeq quantifications for those genes, we cannot produce a quantitative comparison. Fourth, genes classified as pseudogenes in RefSeq may not be pseudogenes in all annotations. As described in Supplementary Data 3, Ensembl considers some of the RefSeq pseudogenes as having a different biotype, such as protein-coding, snRNA, lncRNA. This means that these genes might have had a proper annotation, and not simply a gene start and end coordinates, if RefSeq would be to consider them as a different biotype.

The differences between Ensembl versions also highlight the important question of what needs to be annotated? Adding information to an annotation can have an impact on already existing annotations, as seen here and as previously described^53^. As an example, Ensembl is seen as having more read-through transcripts than RefSeq, and this number is getting bigger. Some of these transcripts have been shown to only be expressed in a cancer cell line, such as the HHLA1-OC90 read-through transcript in teratocarcinoma^54^, while very few are actually found to be expressed more than anecdotally in non-cancerous cells^55^. As soon as a read-through annotation is added, count-based quantifiers will redistribute the counts of the overlapping genes. This should prompt users to move towards more appropriate software, but it also raises questions about the suitability of a ubiquitously used genome annotation, where annotation of rare events might affect the analysis.

Our ICA approach to identify gene group specific biases has proven itself useful, but must be used thoroughly and interpreted by a knowledgeable user. Removing quantification software from the model has produced a technical mode that was unseen in the first model. This means that generating ICA models using exhaustive combinations of inclusion and exclusion of software might reveal more technical biases that were hidden by present technical modes. Including a broader diversity of software would probably also deliver a larger diversity of technical modes. Transcriptome-based software and pseudo-aligners, despite the added difficulty of the combined steps, should be studied considering their growing place in the literature.

We believe that our approach contributes in answering the “what-to-benchmark” question. Our study provided data supporting the concept that the choice of a genome annotation plays an important role in gene quantification. This bias, like the others observed through the ICA for several pipeline steps, is not global, but affects specific gene groups sharing common features. We must emphasize the genome annotations because we believe that, in opposition to alignment and quantification tools, they have not received an appropriate amount of interest with respect to their importance in the definition of the results.

## DATA AVAILABILITY

All codes used for the analyses described in this work are available as a Snakemake workflow (http://gitlabscottgroup.med.usherbrooke.ca/Joel/permutations-ica).

## ACKNOWLEDGEMENT

The authors wish to thank members of their groups for insightful discussions. The authors acknowledge Compute Canada as an outstanding resource for Canadian researchers.

## FUNDING

This work was supported by the Natural Sciences and Engineering Research Council of Canada [NSERC grant RGPIN-2018-05412 to MSS]. JS was supported by Masters scholarships from FRQNT and NSERC. MSS holds a Fonds de Recherche du Québec – Santé (FRQS) Research Scholar Junior 2 Career Award.

## CONFLICT OF INTEREST

None declared.

