## Supplementary Figures and Tables for "Factorial study of the RNA-seq computational workflow identifies biases as technical gene signatures"

### Supplementary materials

*[1] Department of Biochemistry and Functional Genomics, Faculty of Medicine and Health Sciences, Université de Sherbrooke, Sherbrooke, Québec, J1K 2R1, Canada.*

*[2] Department of Chemical & Biotechnological Engineering, Faculty of Engineering, Université de Sherbrooke, Sherbrooke, Québec, J1K 2R1, Canada.*

| **Tissue** | **Sample ID** |
|---|---|
| Colon | ERR315348 |
| Colon | ERR315357 |
| Colon | ERR315400 |
| Colon | ERR315462 |
| Heart | ERR315328 |
| Heart | ERR315331 |
| Heart | ERR315356 |
| Heart | ERR315384 |
| Testis | ERR315351 |
| Testis | ERR315352 |
| Testis | ERR315456 |
| Testis | ERR315492 |
| Thyroid | ERR315397 |
| Thyroid | ERR315412 |
| Thyroid | ERR315422 |
| Thyroid | ERR315428 |

**Supplementary Table 1 | Samples and tissues used in this study**

Samples tissues and ID from the Array-Express E-MTAB-2836 datasets (https://www.ebi.ac.uk/arrayexpress/experiments/E-MTAB-2836/)

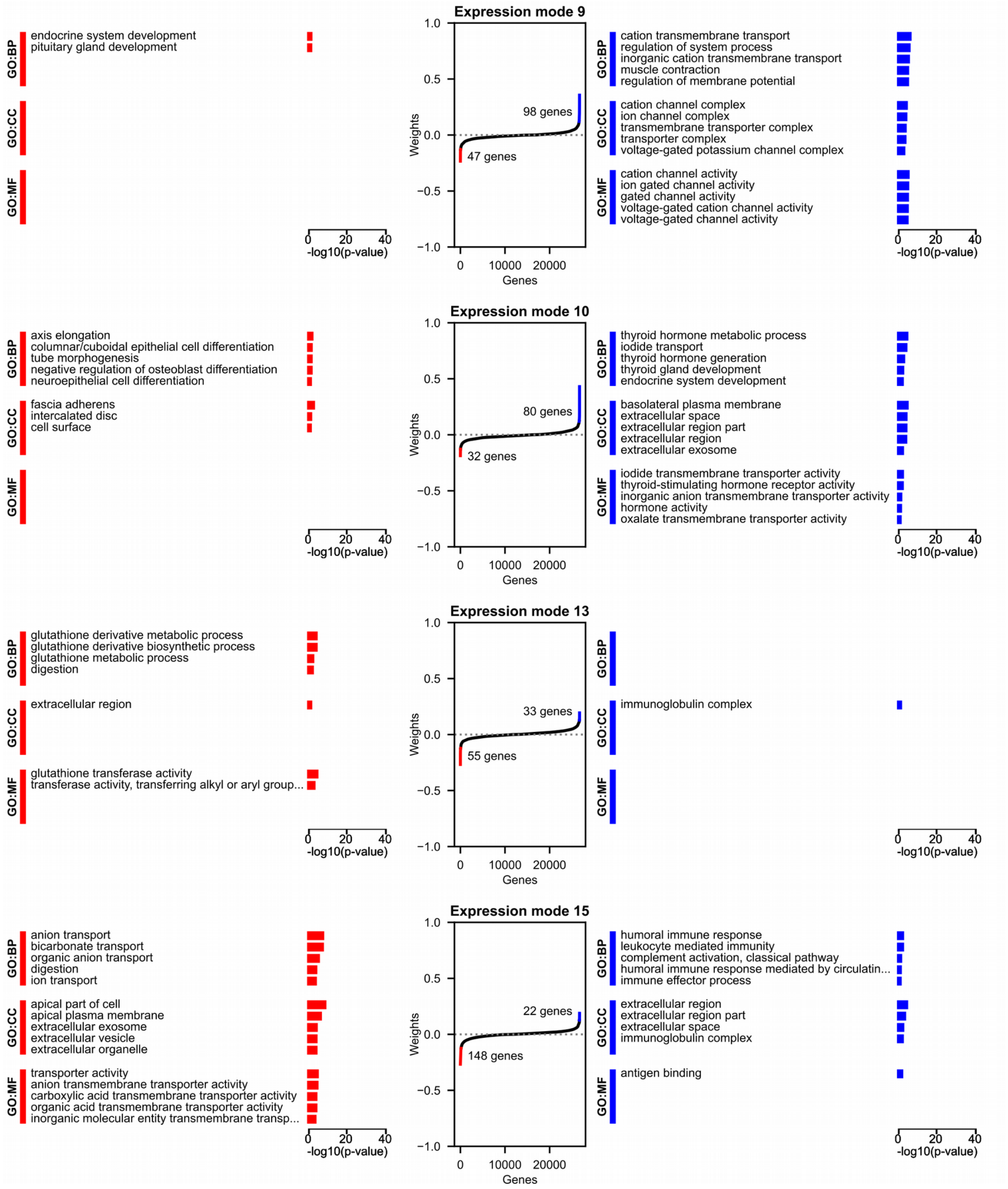


**Supplementary Figure 1 | GO enrichment analysis for biological modes**

The gene weight distributions and the two-sided Gene Ontology (GO) enrichment analysis for the four biological modes presented in Figure 2C are displayed here. For every biological mode, the side with the largest number of genes also has GO terms that are related to the tissue that the biological mode is able to classify.

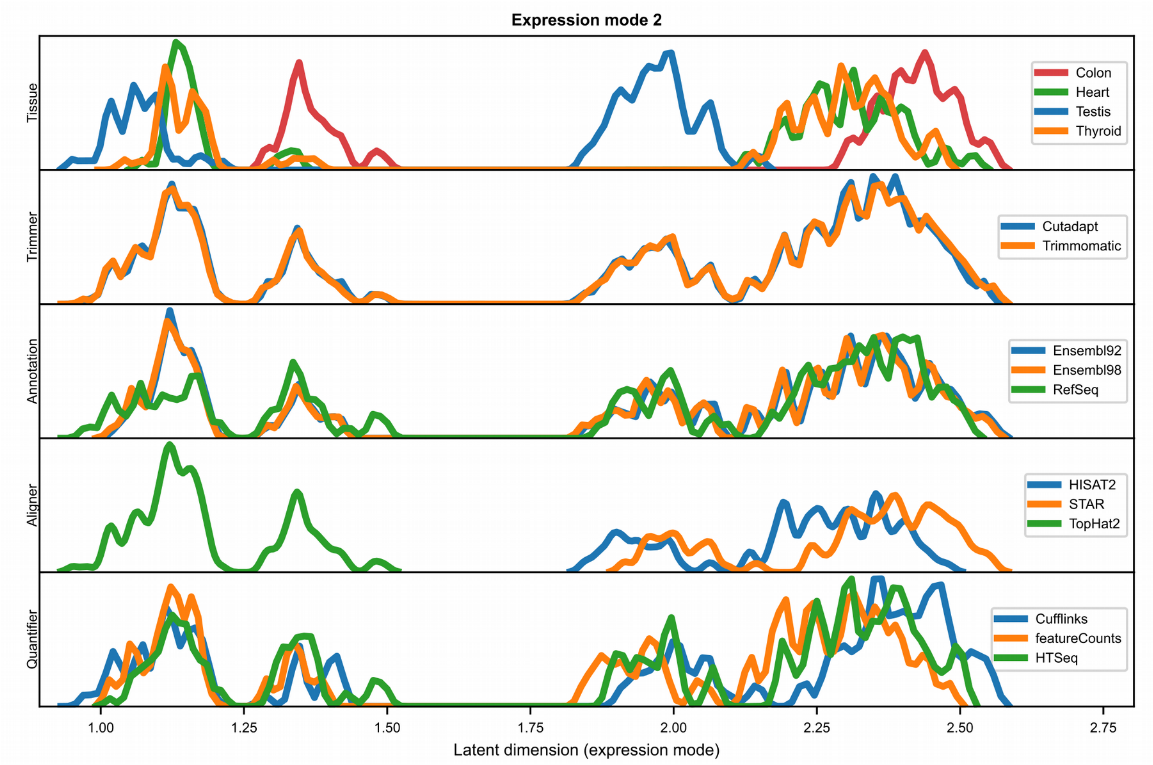


**Supplementary Figure 2 | Projections of expression mode 2**

Distribution of the pipelines along the latent dimension of the expression mode 2 (EM2). The same distribution is shown in five occurrences, one for each pipeline step. Within each occurrence, the pipelines are separated according to the tool used. This representation helps us interpret the information used by the expression mode. For EM2, we can observe that the distribution is mainly driven by the alignment software, since TopHat2 is clustered alone, clearly separated from STAR and HISAT2. We can also see that some of the variability in the shape of the alignment clusters is due to biological variability between tissues, with clusters of colon and testis tissues clustering apart from the other tissues. We might interpret this as if some important genes in defining the alignment clustering have large expression differences in these tissues.

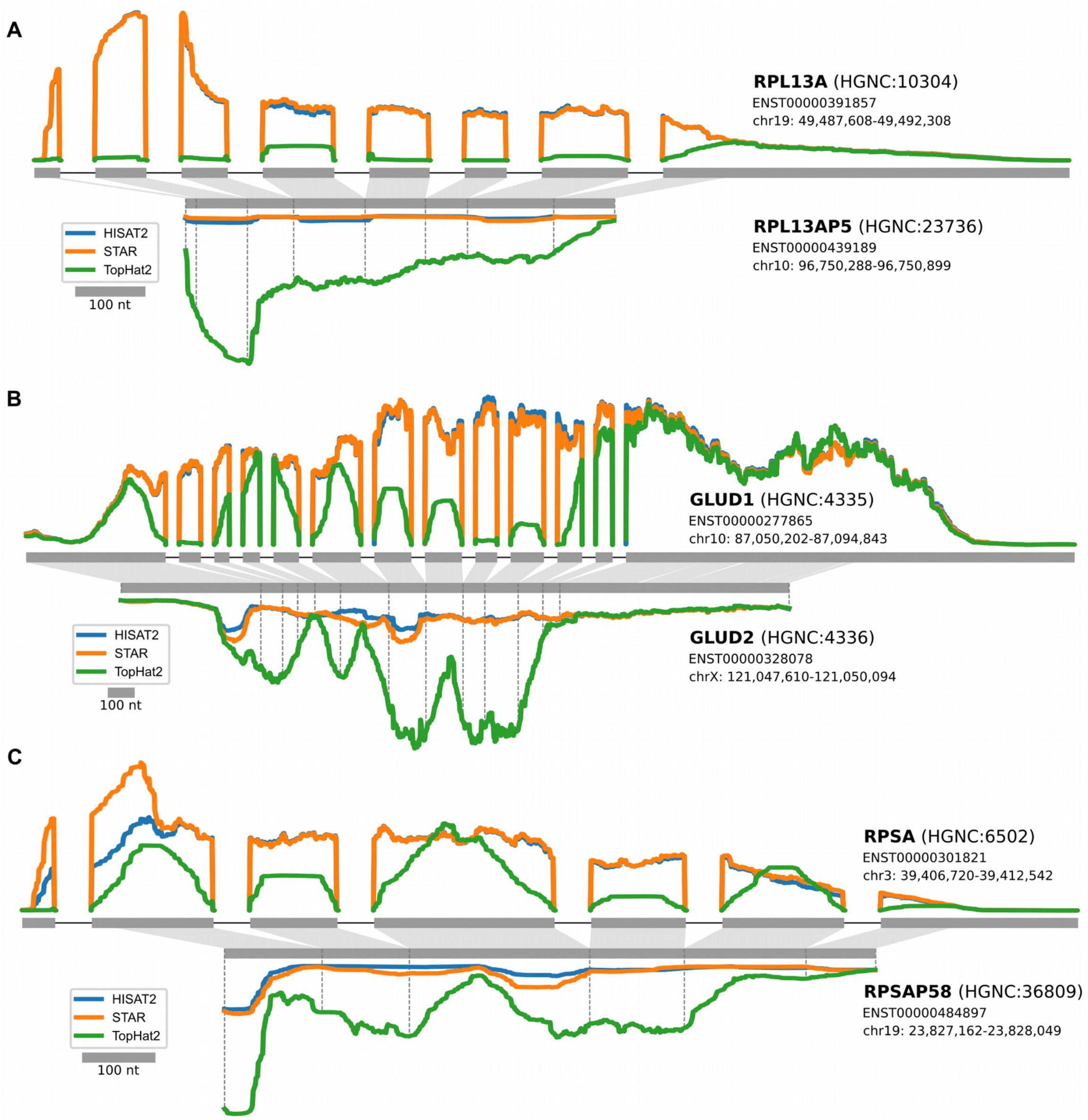


**Supplementary Figure 3 | Gene and pseudogene read profiles**

Comparison of the read profiles along paired genes and pseudogenes for TopHat2, HISAT2 and STAR. This representation is the same as described in Figure 3F.

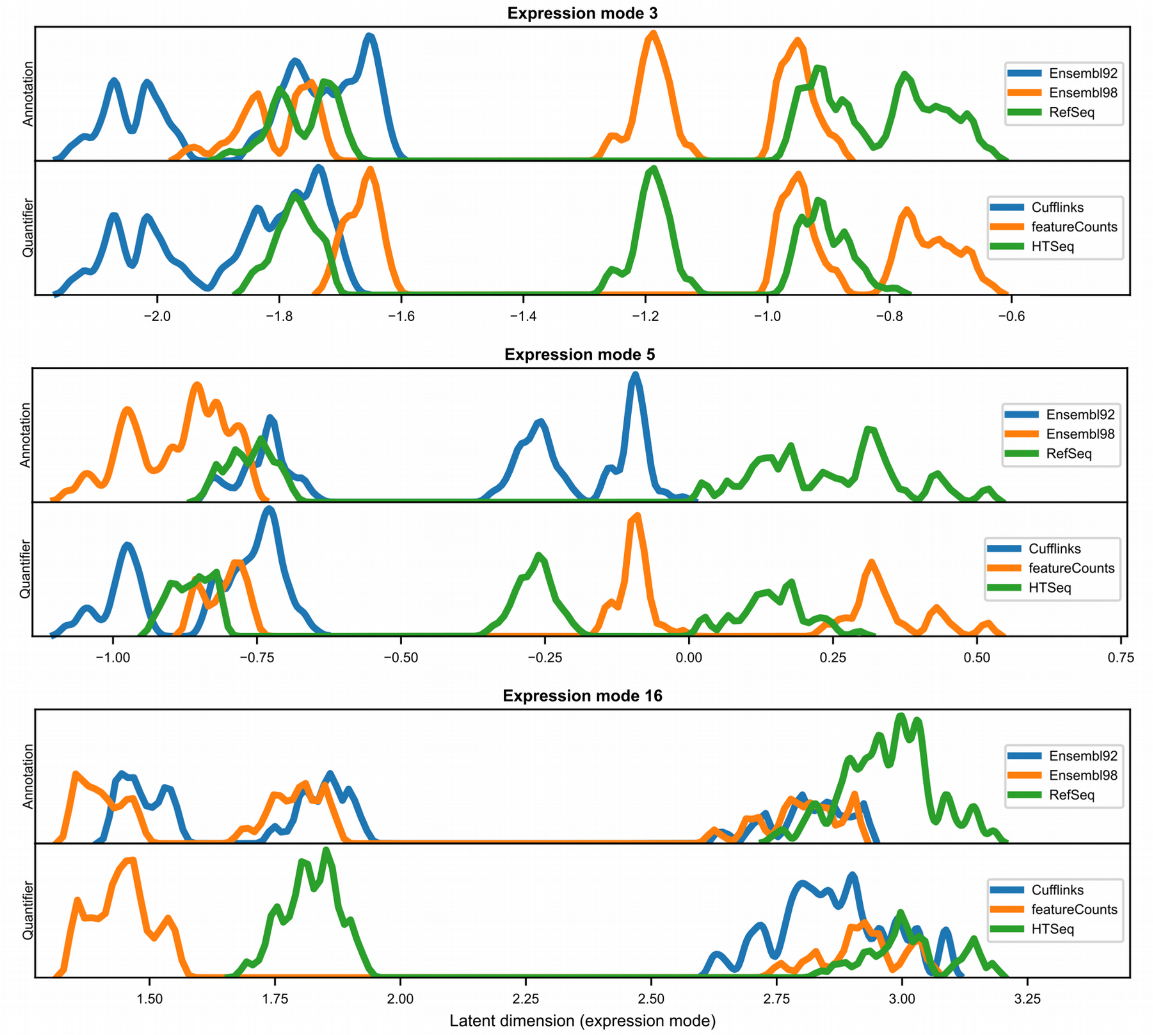


**Supplementary Figure 4 | Projections of annotation and quantifier related technical modes**

Expression modes 3, 5 and 16 are all partly clustered for the genome annotation and the quantifier tools. Distributions of the pipeline along the expression modes are presented for the annotation and the quantifier. In every distribution, Cufflinks and a genome annotation are clustered on one side, whereas the two other quantifiers and annotations are separated between the two sides.

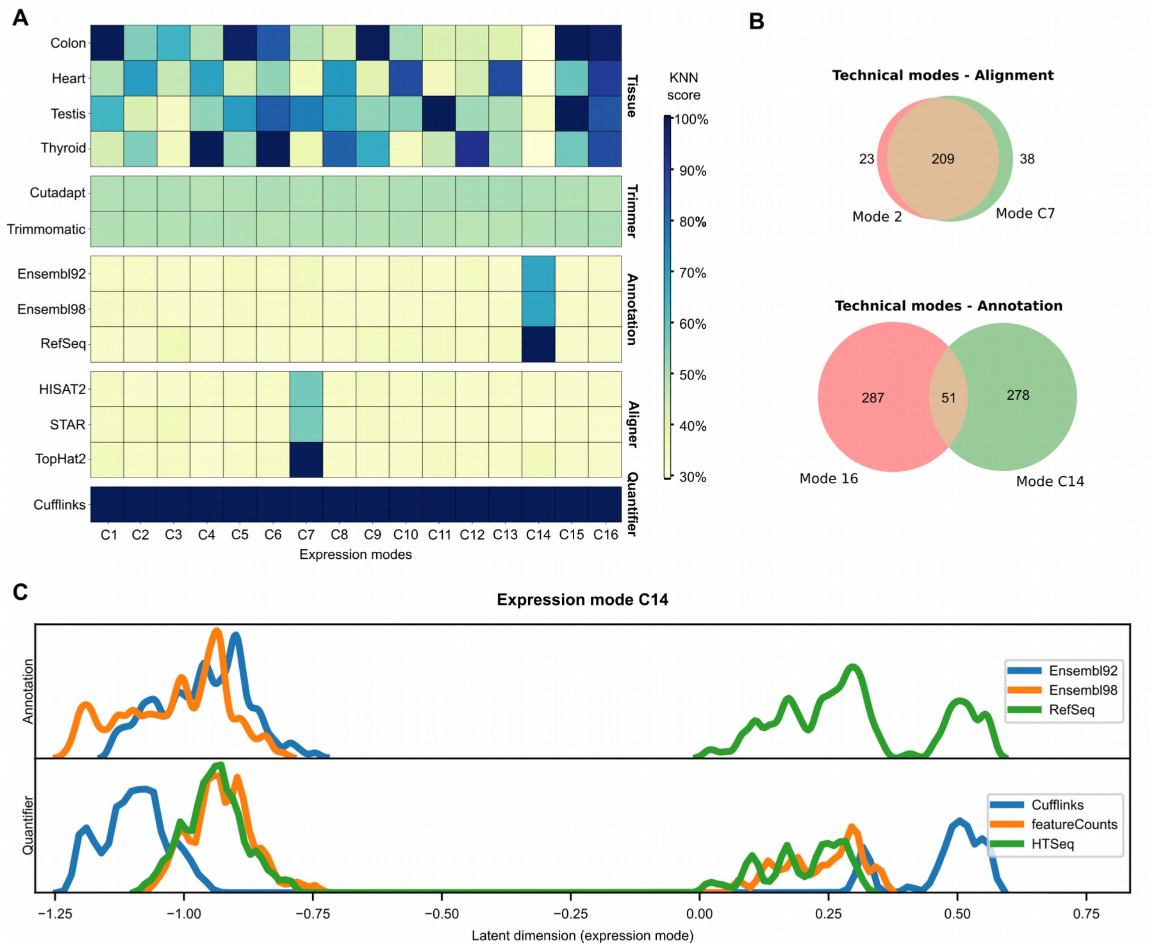


**Supplementary Figure 5 | ICA model for quantification using Cufflinks**

Results and observations of an ICA model that was computed by exclusively using expression datasets that were quantified through Cufflinks. **A** presents the KNN score heatmap for expression mode classification. Only two technical modes, C7 and C14 were detected, respectively linked to aligners and genome annotations. **B** quantifies the overlap of significant genes between technical modes linked to the alignment and annotation from this model to those from the original ICA model. **C** illustrates the distribution of pipelines along C14. In this instance, expression datasets from featureCounts and HTSeq were projected along the EMC14, even if they were not part of the model datasets.

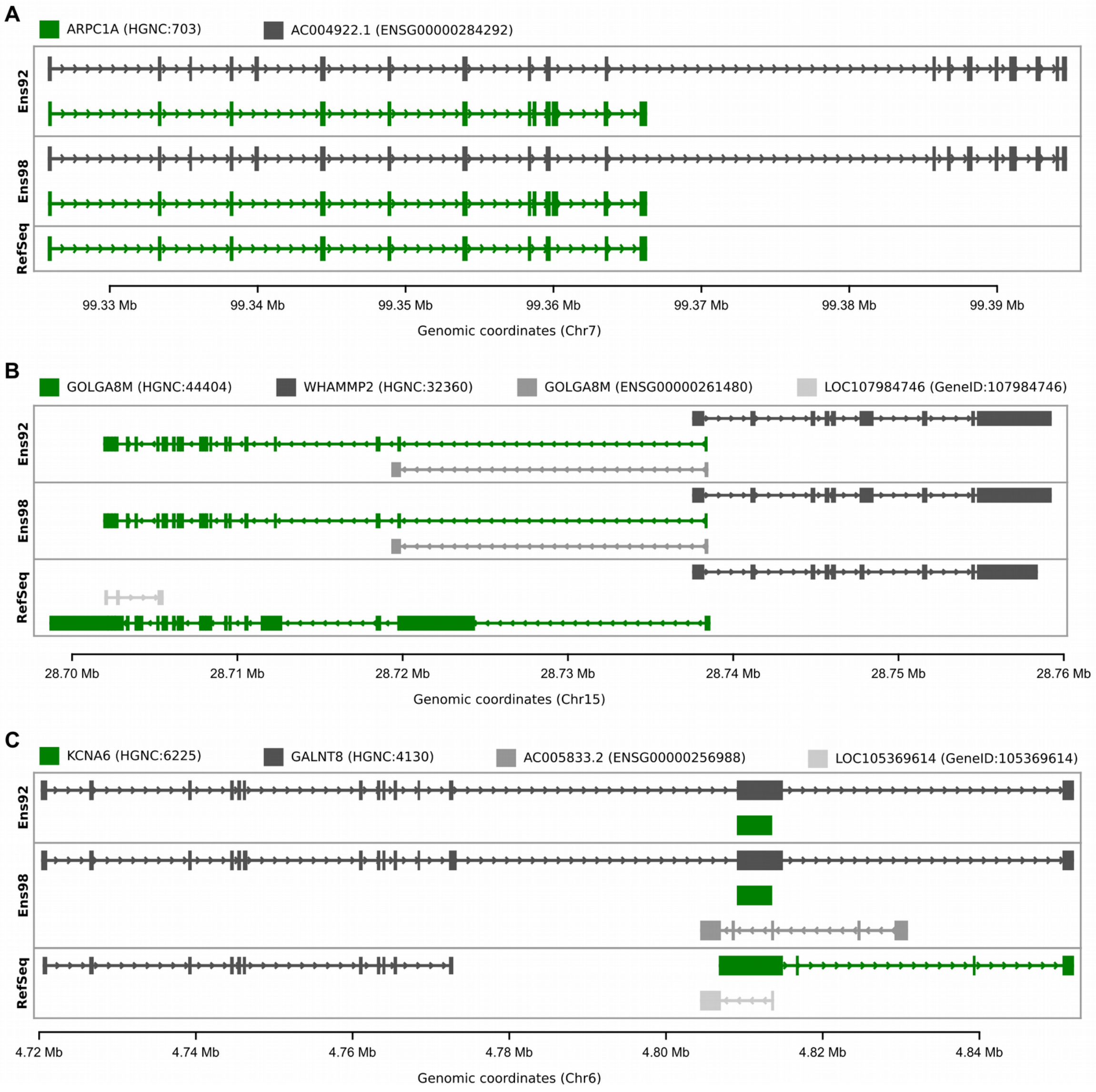


**Supplementary Figure 6 | Ensembl versus RefSeq discriminating genes**

Each gene is summarized as a single entry with a single level of information, where the displayed structure of the gene is a one-dimensional projection, the shadow, of all the different isoforms. Each position in a block is part of an exon in a least one isoform, whereas each position not in block is an intron in all the different isoforms. The plots are centered on a gene of interest, the green gene, and display all the other genes that share at least one chromosomal coordinate in at least one annotation. Each gene of interest is displayed in the three studied genome annotations, for comparison. The genes are identified by their symbol, and the most common identifier (e.g. a HGNC identifier if they possess one, or their genome annotation specific identifier in the second case). The three genes of interest were found in a technical mode related to the classification between Ensembl and RefSeq. ARPCA1 (**A**) is a protein-coding gene found in EM16, GLOGA8M (**B**) is a protein-coding gene found in EMC14 and KCNA6 (**C**) is a protein-coding gene found in both EM16 and EMC14.

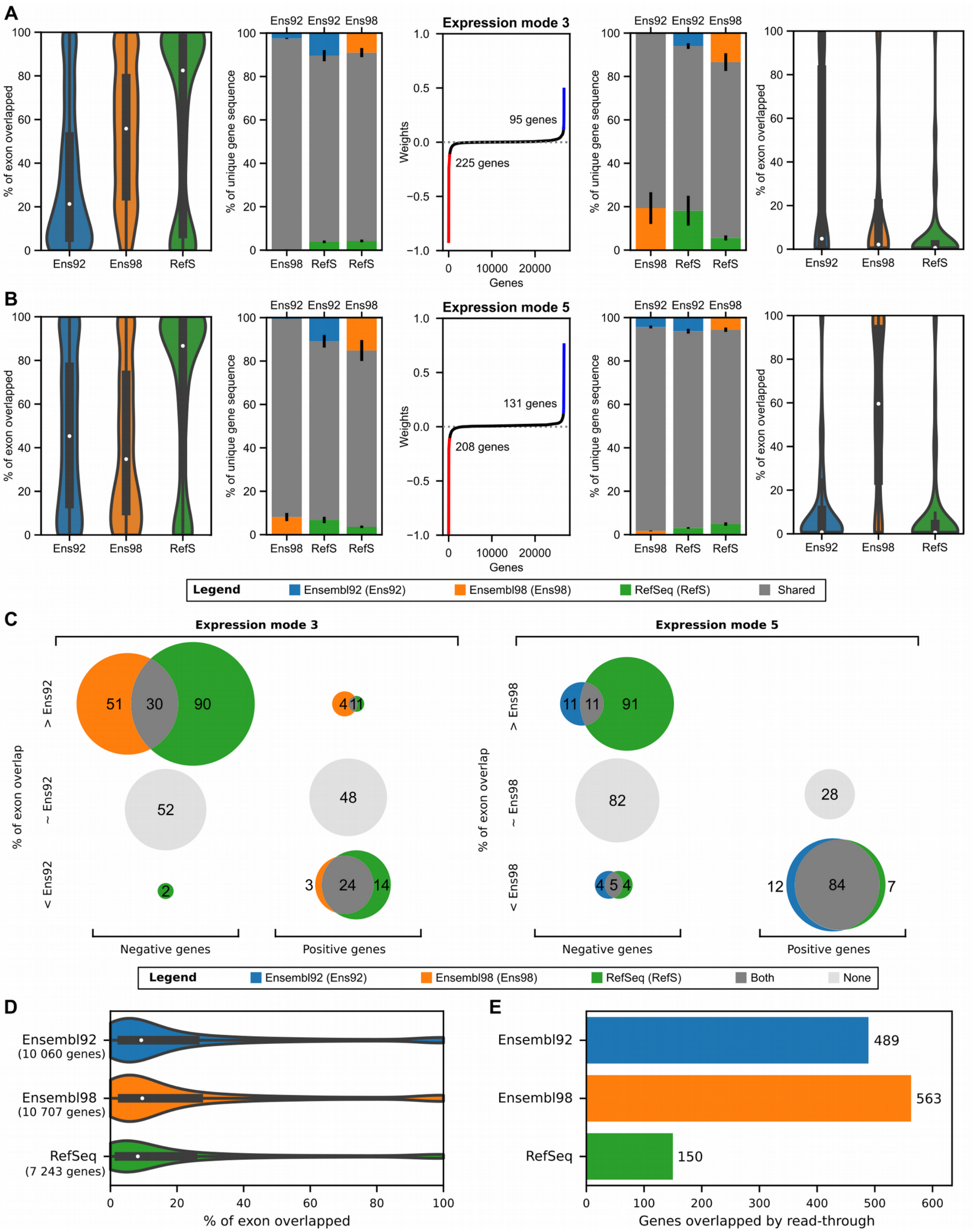


**Supplementary Figure 7 | Evidence for Ensembl 92 versus Ensembl 98 technical modes**

**A** and **B** present the plots described in Figure 4 for EM3 and EM5, highlighting the features discriminating the different genome annotations. **C** classifies the positive and negative genes from EM3 and EM5 according to their percentage of exon overlapped, relative to the genome annotation that is clustered individually. Genes with a score within 10% of the reference annotation score, or with two bigger but opposite scores, were identified as being similar to the reference annotation. Genes with score at least 10% bigger or smaller than the reference annotation were classified as such, colored accordingly to which scores were significantly different. **D** quantifies the percentage of exon overlapped by other genes, either on the sense or antisense strands, for all the 26 713 genes considered in this study. Only the genes with non-null overlap were used to produce the violin plots, and their quantity is presented with the genome annotation names. **E** quantifies the number of read-through genes overlapping at least one gene included in this study.

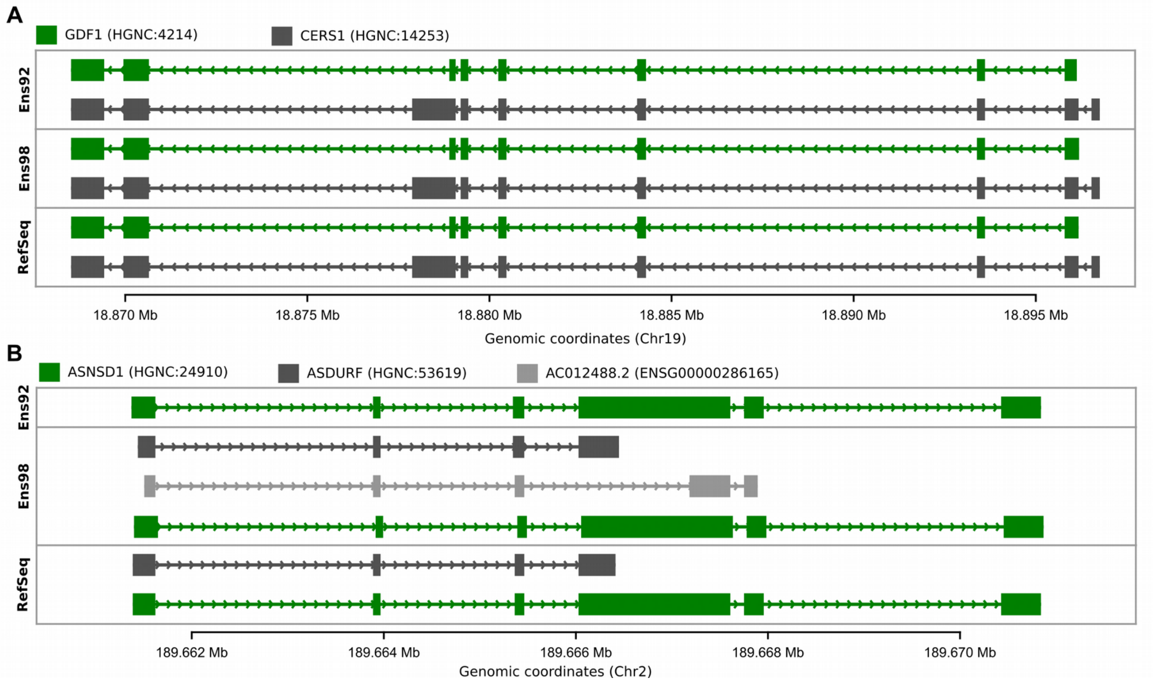


**Supplementary Figure 8 | GDF1 and CERS1 are highly overlapped genes**

**A.** CERS1 and GDF1 are two overlapping genes that share a large proportion of their exons, and that are annotated in the same manner across the three studied annotations. **B.** ASNSD1 transcripts were split into three different genes from Ensembl version 92 to version 98. One of the new genes also has an HGNC ID, and is found in RefSeq. The gene representation is the same as described in Supplementary Figure 5.

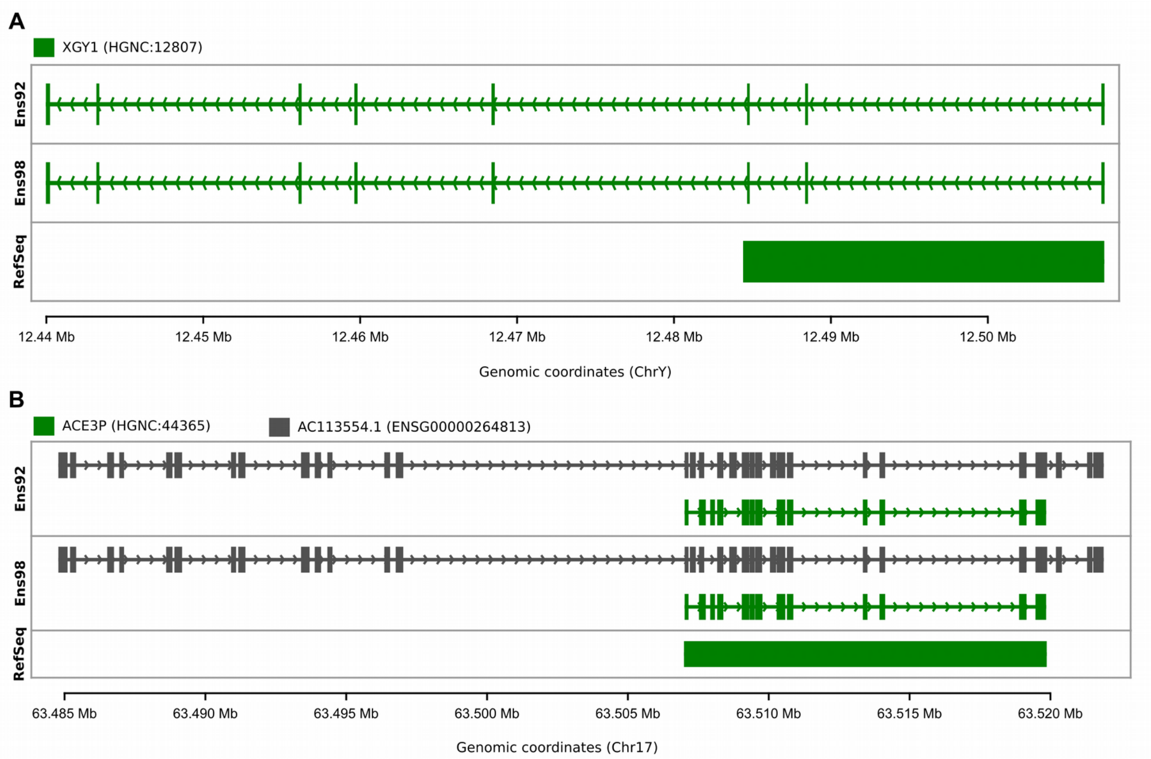


**Supplementary Figure 9 | Example of unprocessed pseudogenes**

A and B represent pseudogenes that are differently annotated in RefSeq and Ensembl. The gene representation is the same as described in Supplementary Figure 5.
